# GWAS and functional studies implicate a role for altered DNA repair in the evolution of drug resistance in *Mycobacterium tuberculosis*

**DOI:** 10.1101/2022.01.04.474954

**Authors:** Saba Naz, Kumar Paritosh, Priyadarshini Sanyal, Sidra Khan, Yogendra Singh, Umesh Varshney, Vinay Kumar Nandicoori

## Abstract

The emergence of drug resistance in *Mycobacterium tuberculosis* (*Mtb*) is alarming and demands in-depth knowledge for timely diagnosis. We performed genome-wide association analysis (GWAS) using 2237 clinical strains of *Mtb* to identify novel genetic factors that evoke drug resistance. In addition to the known direct targets, for the first time, we identified a strong association between the mutations in the DNA repair genes and the multidrug-resistant phenotype. To evaluate the impact of variants identified in the clinical samples in the evolution of drug resistance, we utilized knockouts and complemented strains in *Mycobacterium smegmatis (Msm)* and *Mtb*. Results show that variant mutations abrogated the function of MutY and UvrB. MutY variant showed enhanced survival compared with wild-type (*Rv*) when the *Mtb* strains were subjected to multiple rounds of *ex vivo* antibiotic stress. Notably, in an *in vivo* Guinea pig infection model, the MutY variant outcompeted the wild-type strain. Collectively, we show that novel variant mutations in the DNA repair genes abrogate their function and contribute to better survival under antibiotic/host stress conditions.

## Introduction

The acquisition of drug resistance in *Mtb* has evoked a precarious situation worldwide (1). TB cases where an effective drug regimen is unavailable are inclined towards the development of drug resistance. Resistance to both isoniazid and rifampicin, the first-line drugs, results in multi-drug resistant-TB (MDR-TB). In addition, when the pathogen becomes resistant to fluoroquinolones and one second-line injectable drug (amikacin or kanamycin or capreomycin), it is termed as extremely drug resistant-TB (XDR-TB) (1). Prolonged treatment duration, high drug toxicity, and the expensive drug regimen pose a challenge for treating the MDR and XDR-TB. Besides, the inadequate treatment of drug-resistant TB leads to the augmentation of resistance to other anti-TB drugs, increasing the probability of transmission of these strains in the population (2, 3).

Seven major lineages of *Mtb* are present across the globe, out of which four lineages-Lineage 1-Indo Oceanic (EAI); Lineage 2- Beijing; Lineage 3- Central Asian (CAS) and Lineage 4- Euro-American (4) are prevalent in humans. Although *Mtb* has a lower mutation rate, it gains drug resistance in clinical settings (5). Clinical strains that belong to lineage 2 are more prone to develop drug resistance faster than the lineage 4 strains (6). Acquisition of drug resistance in *Mtb* is majorly attributed to the chromosomal mutations that either modify the antibiotic’s direct target or increase the expression of efflux pumps that helps in decreasing the effective concentration of the drug inside the cell. The expression of drug modifying/degrading enzymes also contributes towards the acquisition of drug resistance (7). Despite well-known mechanisms of drug resistance, in 10-40% of the clinical isolates of *Mtb*, drug resistance cannot be determined by the mutations in the direct targets of antibiotics, implying the presence of hitherto unknown mechanisms that foster the development of resistance in *Mtb* (8). The current knowledge of the mechanisms and biological triggers involved in the evolution of MDR or XDR-TB is inadequate. However, this knowledge is crucial for developing new drug targets and improved diagnosis. Multiple efforts have been made to determine the mechanisms for the emergence of MDR and XDR-TB. While genome-wide association studies (GWAS) identified different genes that abets the emergence of drug resistance but only for a few genes such as *ponA1, prpR, ald*, *glpK,* and the mutation in the *thyA-Rv2765 thyX-hsdS.1* loci are validated (9–12).

In a quest of identifying genetic triggers that aids in the evolution of antibiotic resistance in *Mtb*, we performed GWAS using global data set of 2237 clinical strains that consist of antibiotic susceptible, MDR and XDR. Interestingly, we have identified mutations in the multiple DNA repair genes of *Mtb* that are associated strongly with the MDR phenotype. Functional validation of the identified mutations of DNA repair enzymes revealed that perturbation in the DNA repair mechanisms results in the enhanced survival of strains in the presence of antibiotics *ex vivo* and *in vivo*.

## Results

### GWAS unveils mutations in the DNA repair genes

To identify the genetic determinants contributing to the development of antibiotic resistance in *Mtb*, we performed genome-wide association analysis using whole-genome sequences of clinical strains from 9 published studies. After the quality filtering of raw reads, our dataset had 2773 clinical strains from 9 different countries and four lineages; Lineage 1-Indo Oceanic (EAI); Lineage 2- Beijing; Lineage 3- Central Asian (CAS) and Lineage 4- Euro-American (Fig1a & Table S1) (10, 11, 13–19). The dataset majorly represented Lineage 2 and Lineage 4 isolates, which are predominant across the globe. Strains were categorized as susceptible, mono-drug resistant, MDR, Poly-Drug resistant (Poly-DR), and pre-XDR based on computational prediction and phenotypes reported in the previous studies (Table S1) (20). We identified ∼160,000 Single Nucleotide Polymorphism (SNPs) and indels after mapping the short reads on the reference *Rv* genome. The total number of SNPs observed for susceptible, mono-DR, MDR, or pre-XDR strains were comparable, suggesting no genetic drift during the evolution of antibiotic resistance (Fig 1b).

**Figure 1.**
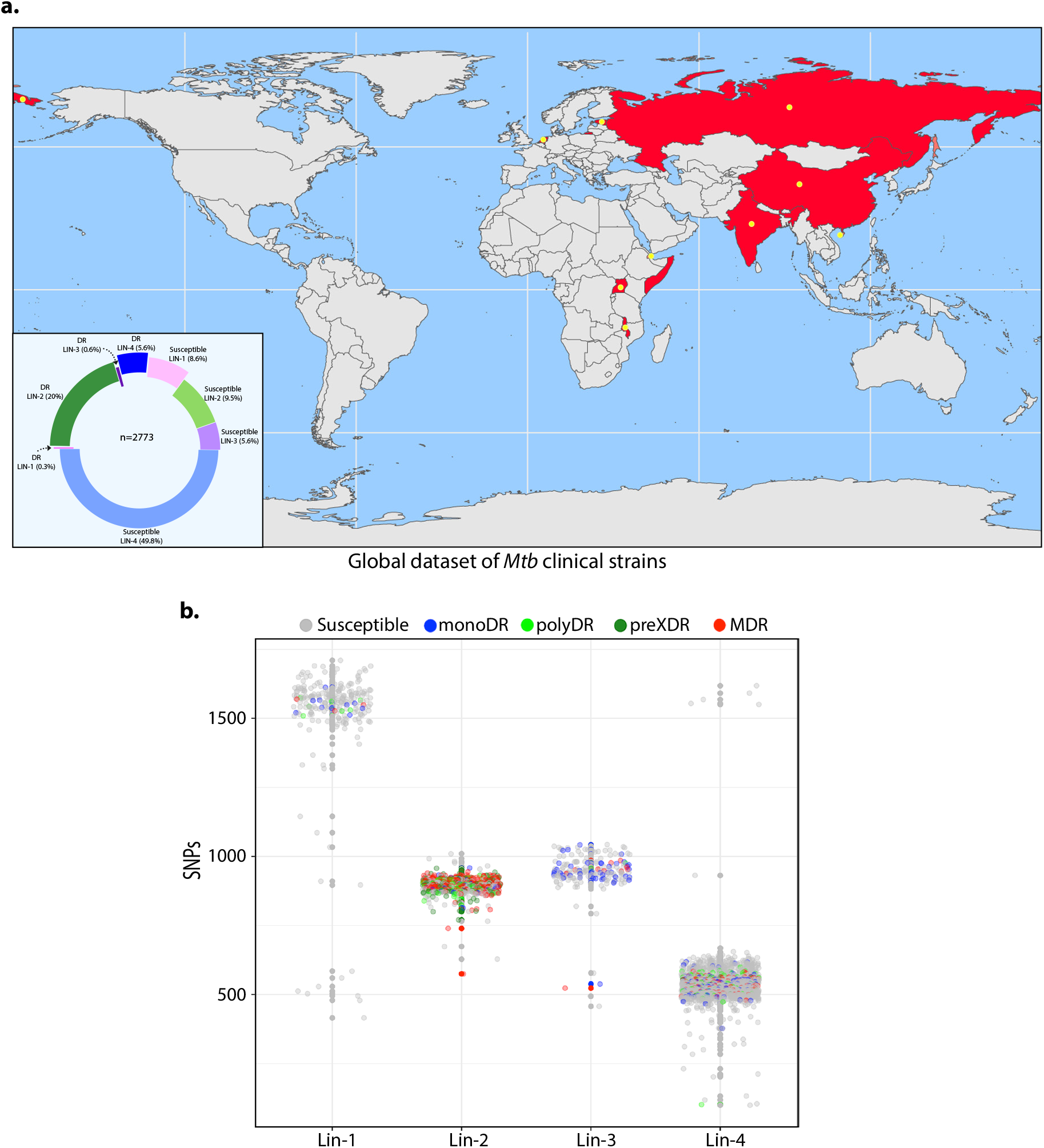
GWAS unveils mutations in the DNA repair genes. **a)** Geographical distribution of 2773 clinical strains of *Mtb*. Donut plot represents the proportion of susceptible and drug-resistant (DR) strains in each lineage. DR includes mono-DR, poly-DR, MDR and pre-XDR. A detailed breakup of distribution is given in Table S1. **b)** Dot-plot showing the number of SNPs identified in each strain. Different colored dots indicate the drug resistance phenotype of strain.

We designed our strategy based on the reasoning that the probability of finding the genetic determinants that contribute to drug resistance would be higher in the strains that are resistant to more than two antibiotics. Thus, we performed GWAS using 1815 drug-susceptible strains and 422 MDR/XDR strains (Fig S1 & Table S2). We employed a genome association and prediction integrated tool (GAPIT) software with a compressed mix linear model under the R environment for GWAS analysis. GAPIT employs population structure and relative kinship matrix during association analysis. The false discovery rate (FDR) adjusted p-values in the GAPIT software are highly stringent as it corrects the effects of each marker based on the population structure (Q) as well as kinship (K) values (21, 22). Therefore the probability of identifying the false-positive SNPs is very low.

After setting the adjusted p-value cut-off at 10^-5^, we identified 188 mutations, including 24 intergenic regions that correlated with the multi-drug resistance (Table S3-S6). The effect of identified SNPs on the development of MDR/XDR reveals positive or negative contributions (Fig 2a). As anticipated, we have identified known first and second-line drug resistance target genes (Fig 2b & Table 1). Although we identified multiple mutations in *rpoB*, only p.Leu452Val and p.Val496Meth were above the cut-off. Notably, mutations in the *rrs, katG* (p.Ser315Thr)*, embB* (p.Gly406Ser)*, pncA*1(p.His71Arg), *gyrA* (p.Ala90Val), and recently reported genetic determinants such as *folC* (p.Ser150Gly), and *pks* were part of the 164 genes, validating our approach (Fig 2b & Table 1 & Table S4 & Table S6). Besides, we identified known the compensatory mutations in the *fabG1* upstream region, *eis-Rv2417c* and *oxyR-ahpC* loci (23) (Table S6).

**Figure 2.**
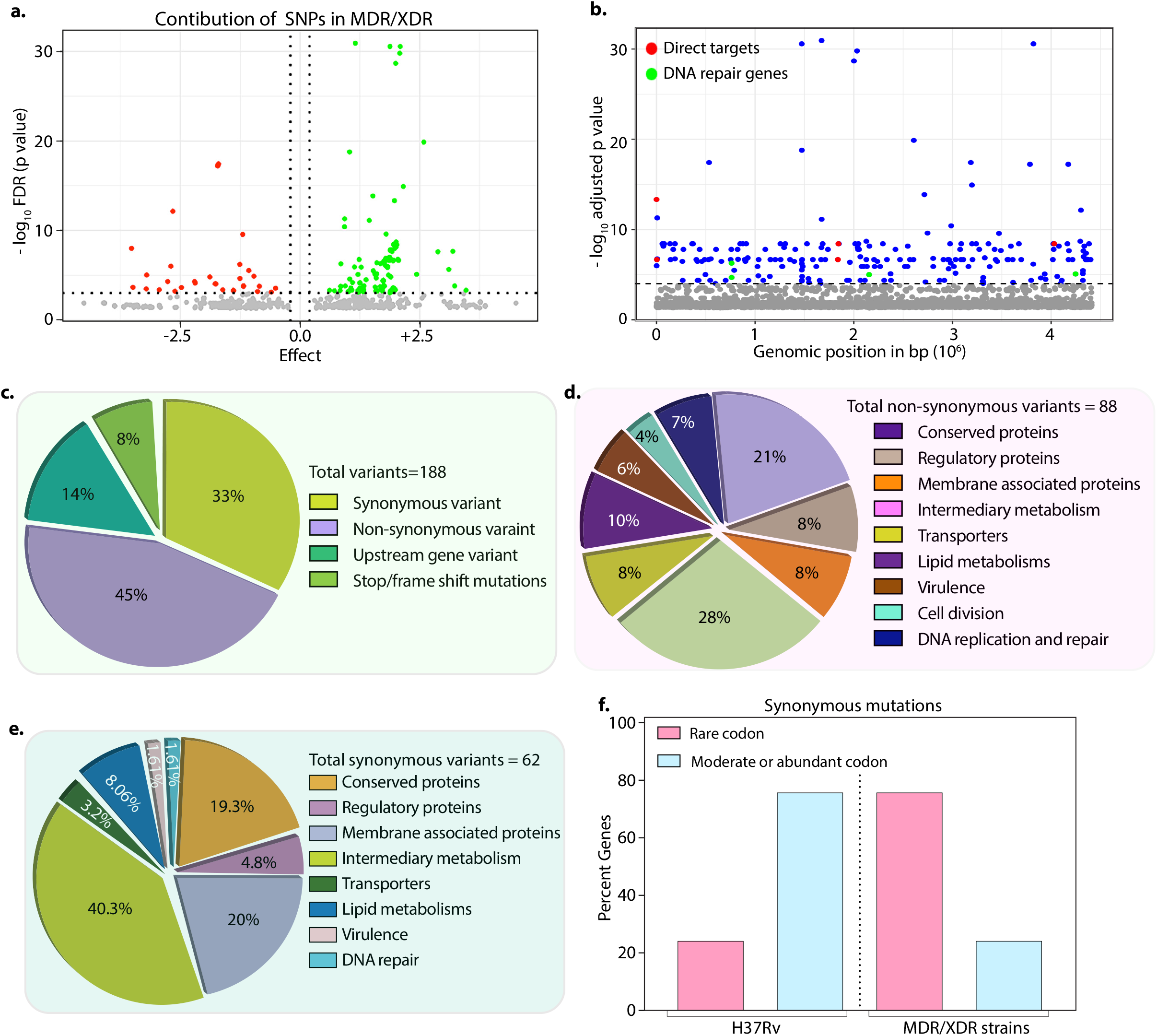
Drug resistant strains carry mutations in the DNA repair genes. **a)** Volcano plot represents the effect of identified SNPs on the development of MDR/XDR-TB. Positive effect (green dots) shows the identified SNPs would aid in MDR/XDR development. The negative effect (red dots) shows that the SNPs would restrain the development of MDR/XDR. **b)** Manhattan plot representing the association between the genes and drug resistance phenotype. A total of 188 genes that include intergenic regions were identified above the 10^-5^ cut-off value through association studies. Blue dots represent mutation in the lipid metabolism, membrane proteins, intermediary metabolism genes, and others. Green dots represent mutation in the direct targets for the first- and second-line antibiotics. Red dots represent mutations associated with the DNA repair genes. A detailed list of associated genes is provided in Table S3 and S4. **c-e)** Pie chart represents the total (c), non-synonymous (d) and synonymous (e) SNPs identified in the genes that belong to different categories. **f)** Bar plot represents the percentage of synonymous mutations in the genes that resulted in abundant/moderate codon usage to the rare codon compared to H37Rv.

**Table 1.**
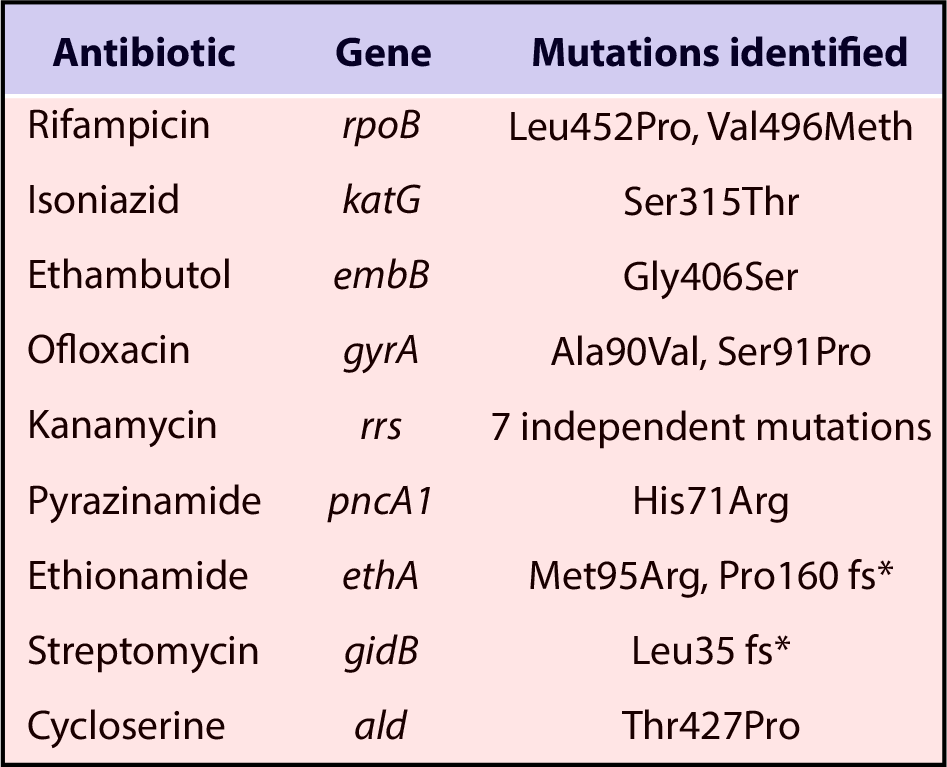
Mutations identified in the direct targets of antibiotic in GWAS analysis.

Among the 188 genes, ∼45% of mutations resulted in non-synonymous changes, whereas ∼33% resulted in synonymous changes, 14% were identified in the upstream gene variant, and 8% possess stop/frameshift mutations (Fig 2c-2e & Table S3-S6). While non-synonymous or frameshift mutations most likely affect the protein function, intergenic region mutations may impact the expression of the gene. For example, we have identified frameshift mutation in *rv2333c* (adjusted p-value 1.32×10^-20^), encoding for a transporter, and a non-synonymous mutation p.Pro561Ser in *rv1250* (adjusted p-value 3.4×10^-7^), known to be differentially expressed in MDR patients, to be strongly associated with the MDR phenotype (Table 3)(24). Besides, we identified mutations in genes involved in the lipid metabolism, intermediary metabolism and respiration, membrane transporters, cell wall and cell processes, membrane-associated proteins, and others (https://mycobrowser.epfl.ch/)(Fig2c-e). On the other hand, synonymous change may alter the mRNA stability or stall the translation process by changing an abundant codon to a rare codon (25–28). Analysis of the synonymous mutations for the codon bias revealed that in 50% of cases, codons are converted moderate/abundant to rare codon (compare 75% in MDR/XDR with 25% in H37Rv) (Fig 2f, Fig S8a & Table S7). This result is in accord with the studies published in other bacteria, where a synonymous mutation impacts mRNA stability (26–29).

In addition to the mutations described above, we identified novel mutations in base excision repair (BER), nucleotide excision repair (NER), and homologous recombination (HR) pathway genes, *mutY, uvrA, uvrB,* and *recF* that are strongly associated with the MDR and XDR-TB (Fig 2b & Table 2). Mutations in the DNA repair pathway genes could contribute towards the selection and evolution of antibiotic resistance (Table 2) (Fig S6). Analysis of the distribution frequency of mutations in the DNA repair genes demonstrated the distribution of mutations specifically in MDR, PDR, and XDR strains (Fig S7a-d). Collectively, GWAS data presented above-identified mutations in the direct targets and yet uncharacterized variants, including novel mutations in DNA repair genes.

**Table 2.** Mutations in DNA repair and replication genes associated with MDR phenotype.

| Gene | Amino acid change | Wild type | Mutated | FDR-adjusted p-value |
| --- | --- | --- | --- | --- |
| *dnaA* | Arg233Gln | G | A | 5.16E-12 |
| *mutY* | Arg262Gln | G | A | 3.83E-09 |
| *uvrB* | Ala524Val | C | T | 2.15E-07 |
| *uvrA* | Gln135Lys | C | A | 3.83E-09 |
| *recF* | Gly269Gly | G | T | 2.15E-07 |

**Table 3.**
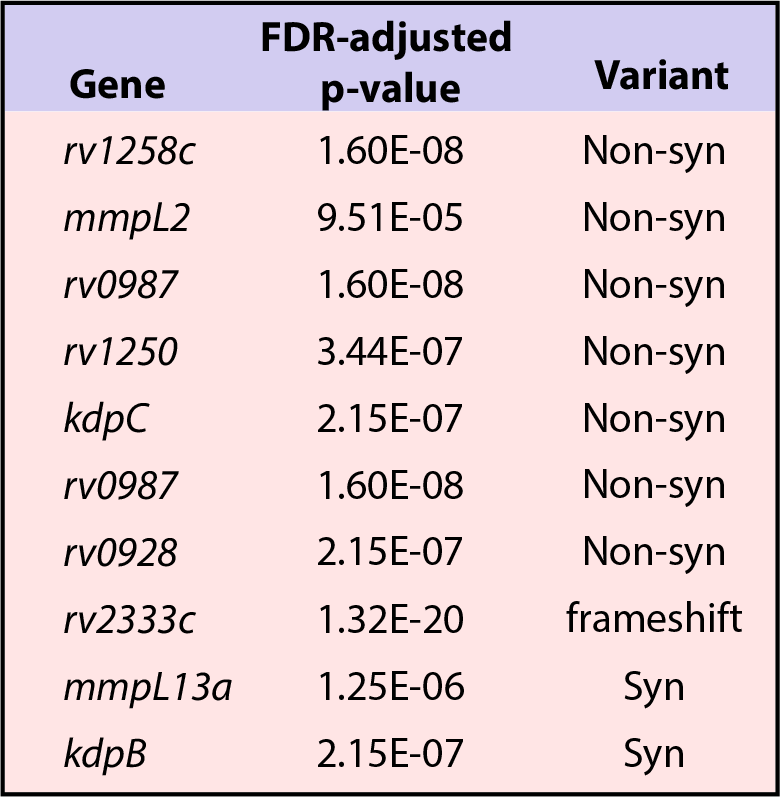
Transporter associated with MDR phenotype.

### Mutations in DNA repair genes result in the loss of function

WGS sequencing of *Mtb* revealed the presence of DNA repair pathways such as BER, NER, HR, Non-Homologous end joining (NHEJ), and Single Strand Annealing (SSA) that guard its genomic integrity (30). MutY is a 302 amino acid (aa) long adenosine DNA glycosylase encoded by *rv3589*. Oxidative damage to the DNA results in the formation of 7,8-dihydro 8-oxoguanine (8-oxoG) that can pair with guanine (G) or adenine (A). If left unrepaired, this would lead to C→G or C→A mutations in the genome (Fig 3a) (31). The GWAS analysis identified **Arg262Gln** mutation at the C-terminal region of the MutY. To decipher the biological role of the identified variant, we cloned *Mtb mutY* and performed site-directed mutagenesis to generate the mutant allele. Wild type and mutant mutY genes subcloned in an integrative *Mtb* shuttle vector. Constructs were electroporated into *Mycobacterium smegmatis mutY* mutant strain (*msm*Δ*mutY*) to generate *msm*Δ*mutY::mutY,* and *msm*Δ*mutY::mutY-*R262Q strains.

**Figure 3.**
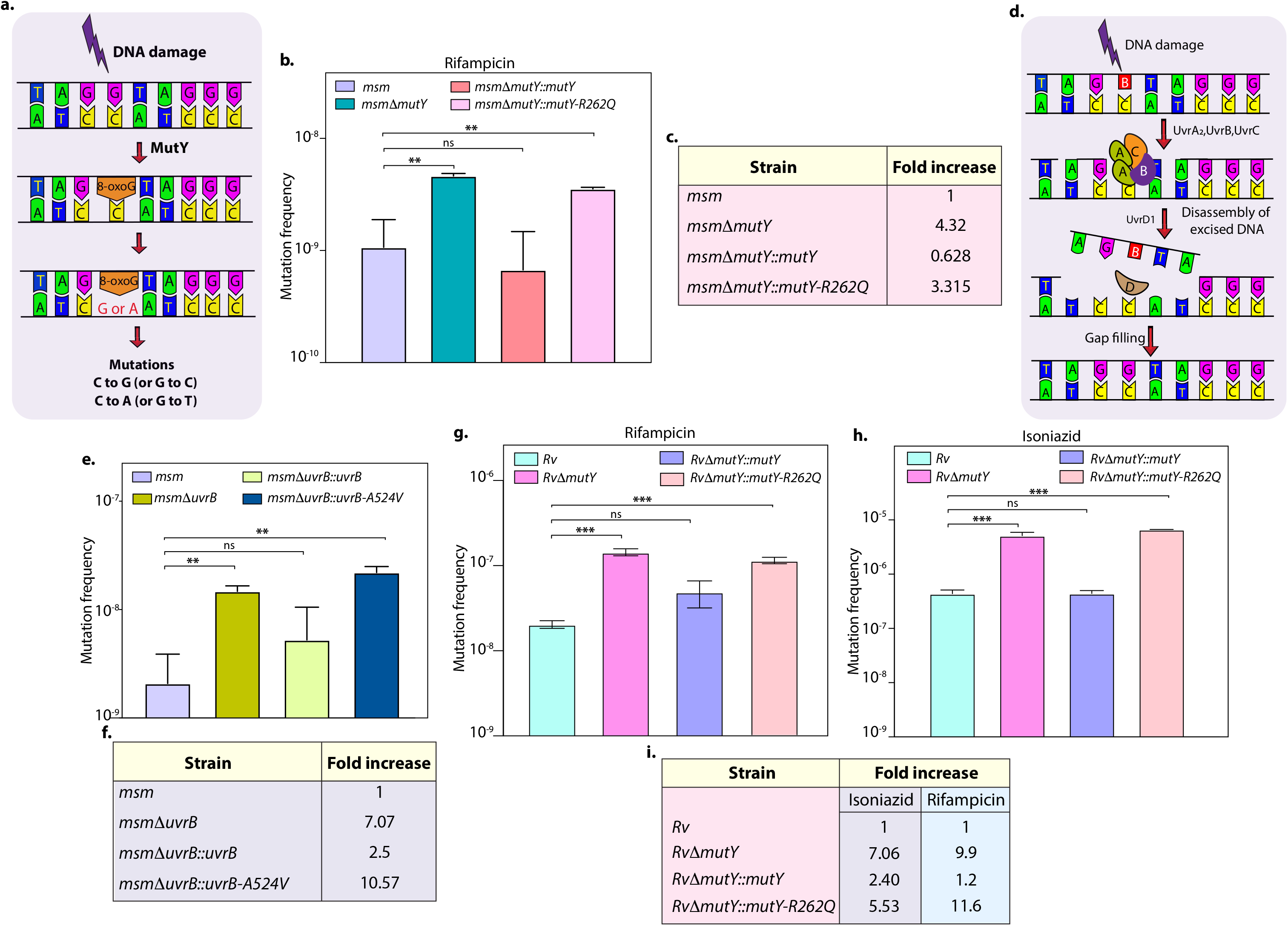
Variants identified in DNA repair genes abrogate their function. **a)** A schematic representation of the BER pathway that operates in mycobacteria. DNA damage results in the generation of 8-oxo-G, which would cause mutations in the genome if left unrepaired. **b)** Mutation frequency was calculated using *msm, msm*Δ*mutY, msm*Δ*mutY::mutY, msm* Δ*mutY::mutY-R262Q*. **c)** Fold increase in the mutation frequency with respect to wild type *msm*. **d)** A schematic representation of the NER pathway showing the recognition and initiation of repair by UvrA-UvrB and UvrC. **e)** Mutation frequency of *msm, msmΔuvrB, msmΔuvrB::uvrB* and *msmΔuvrB::uvrB-A524V*. **f)** Fold increase in the mutation frequency with respect to wild type *msm*. **g & h)** Mutation frequency was calculated for different strains in the presence of rifampicin (g) or isoniazid (h). **i)** Table showing the fold increase in the mutation frequency in comparison with wild-type *Rv*. Two biologically independent experiments with each experiment in technical triplicates was performed. Data represent mean and standard deviation. Statistical analysis (two-way ANOVA) was performed using Graph pad prism software. *** p<0.0001, **p<0.001 and *p<0.01.

We performed mutation frequency analysis to evaluate the impact of Arg262Gln mutation abrogating its DNA repair function. In accordance with the published data, deletion resulted in 4.32 fold increase in the mutation frequency (Fig 3c) (31). While complementation with wild-type *mutY* restored the mutation frequency, complementation with *mutY-R262Q* failed to do so (Fig 3b,c). Next, we investigated the role of mutation identified in NER pathway gene UvrB. UvrA, UvrB, and UvrC recognize and initiate the NER pathway upon DNA damage. UvrB, a 698 aa long DNA helicase, encoded by *rv1633* plays a pivotal role in the NER pathway by interacting with the UvrA and UvrC (Fig 3d) (32). UvrB harbors N and C-terminal helicase domain, interaction domain, YAD/RRR motif, and UVR domain. Identified UvrB variant **Ala524Val** mapped to the C-terminal helicase domain (33). To evaluate the functional significance, *Mtb uvrB* and *uvrB-A524V* genes were cloned in an integrative vector. The absence of uvrB led to higher mutation frequency, which could be restored upon complementation with the wild type, but not with the variant (Fig 3e,f).

Subsequently, we sought to extend our investigations to *Mtb*. Towards this, we generated the gene replacement mutant of *mutY* in laboratory strain H37Rv, wherein the *mutY* at native loci was disrupted with hygromycin resistance cassette. Replacement at the native loci was confirmed by performing multiple PCRs (Fig S8b-c). We determined the mutation frequency in the presence of rifampicin and isoniazid (Fig 3g-i). The fold increase in the mutation frequency relative to *Rv* for *Rv*Δ*mutY*, *Rv*Δ*mutY::mutY*, and *Rv*Δ*mutY::mutY*-R262Q, were 7.06, 2.04, 5.53 in the presence of isoniazid and 9.9, 1.2, 11.6 in the presence of rifampicin (Fig 3g-i). Together the data suggested that variants of *mutY* and *uvrB* identified in the GWAS compromised their function.

### The variant of *mutY* confers survival advantage *ex vivo*

To ascertain the role of the identified variant of *mutY*, we evaluated the survival of *Rv, Rv*Δ*mutY*, *Rv*Δ*mutY::mutY,* and *Rv*Δ*mutY::mutY-*R262Q in the peritoneal macrophages. We did not observe any difference in the survival of *Rv*Δ*mutY* or *Rv*Δ*mutY::mutY-*R262Q when compared with *Rv* and *Rv*Δ*mutY::mutY* (Fig S9a-b). We hypothesized that the evolution of strain to become antibiotic-resistant required continued antibiotic and host-directed stress. Therefore, we infected peritoneal macrophages with *Rv, Rv*Δ*mutY*, *Rv*Δ*mutY::mutY,* and *Rv*Δ*mutY::mutY-*R262Q, in the absence or presence of the antibiotics. The bacteria recovered were cultured *in vitro* for 5 days and used for the next round of infection. The process was repeated for three rounds, and CFUs were enumerated at 4 and 96 h post-infection (p.i) during the fourth round of infection (Fig 4a). CFU’s obtained at 4 h p.i showed equal load. *Rv*Δ*mutY* and *Rv*Δ*mutY::mutY-*R262Q exhibited better survival in the absence of antibiotics compared with *Rv* and *Rv*Δ*mutY::mutY* (Fig 4b-e). There was no additional advantage compared with untreated in the presence of isoniazid (Fig 4f). However, we observed a log-fold advantage for *Rv*Δ*mutY* and *Rv*Δ*mutY::mutY-*R262Q compared with Rv or *RvΔmutY::mutY* in the presence of rifampicin or ciprofloxacin (Fig 4f). Results suggest that antibiotic pressure in the host drives the acquisition of mutations, resulting in the better survival of the strains.

**Figure 4.**
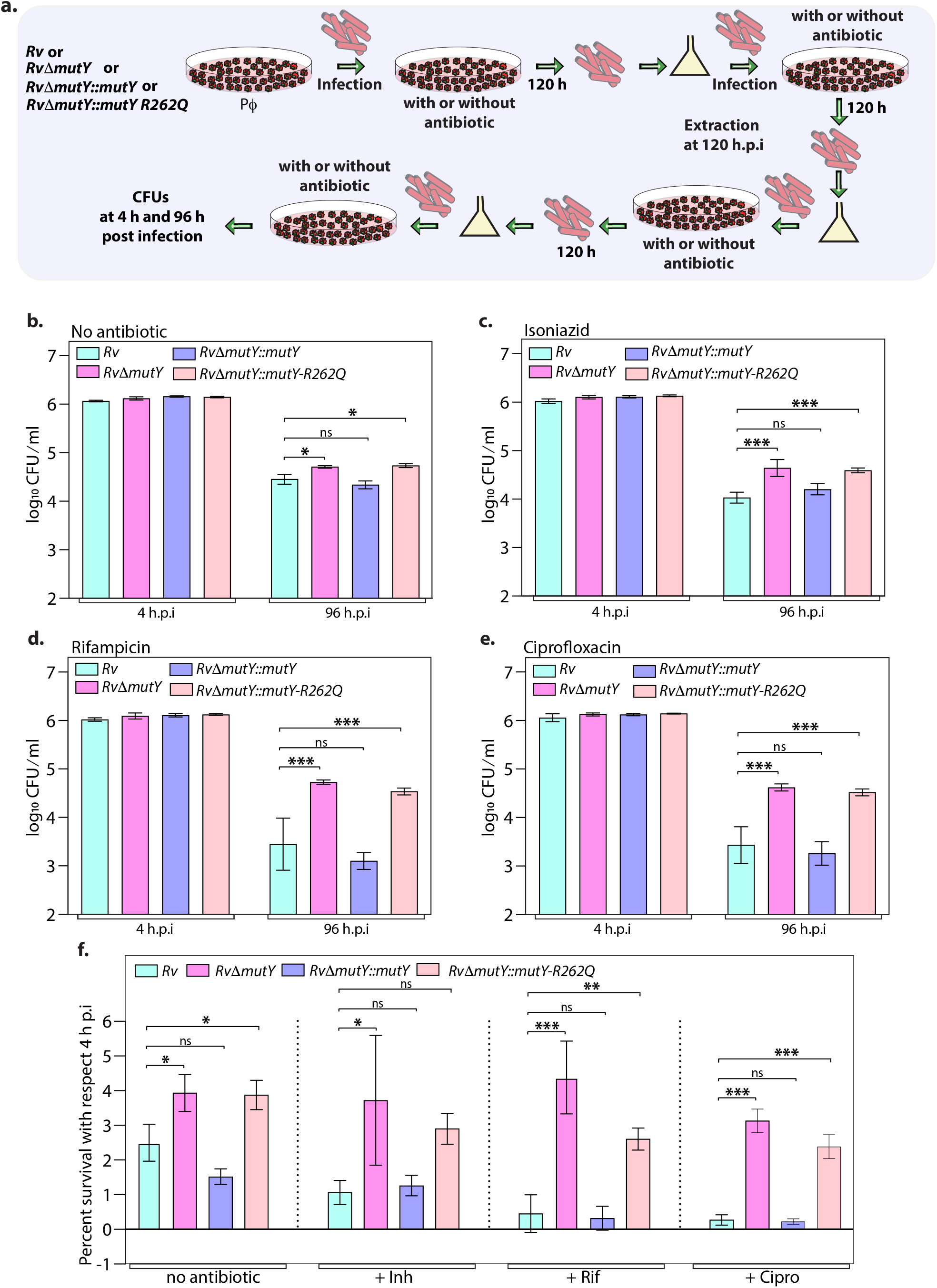
Mutations in the DNA repair genes provide a survival advantage in the presence of antibiotics. **a)** A schematic representing the *ex vivo* infection experiment in the presence and absence of different antibiotics. **b-e)** Survival of the strains in the peritoneal macrophages at 4 and 96 h p.i. without and with antibiotics (isoniazid or rifampicin or ciprofloxacin). **f)** Percent survival with respect to day 4 h p.i was determined for each strain without and with antibiotics (isoniazid or rifampicin or ciprofloxacin). Two biologically independent experiments were performed for determining mutation frequency. Each experiment was performed in technical triplicates. Data represents one of the two biological experiments. Data represents mean and standard deviation. Statistical analysis (two-way ANOVA) was performed using Graph pad prism software. *** p<0.0001, **p<0.001 and *p<0.01.

### Competition experiment ex vivo reveal MutY variant confers survival advantage

Results above suggested that *mutY* variant impacts survival advantage when subjected to antibiotic selection, likely due to its ability to accumulate mutations. We reasoned that, if this is indeed the case, the *mutY* variant may outcompete the wild type *Rv*, when both the strains are present together. To test this hypothesis, we infected peritoneal macrophages with a combination *Rv+Rv*Δ*mutY* or *Rv+Rv*Δ*mutY::mutY* or *Rv+Rv*Δ*mutY::mutY-*R262Q. 96 h p.i cells were lysed and CFUs were enumerated on plain 7H11 or Kanamycin (*Rv*) or Hygromycin (*Rv*Δ*mutY, Rv*Δ*mutY::mutY, Rv*Δ*mutY::mutY-*R262Q) to evaluate the survival rates (Fig S9c-d). The percent survival was calculated as CFUs obtained on Kanamycin or Hygromycin / CFUs on Kanamycin+Hygromycin. It is apparent from the data that the survival rates of competing strains were comparable, suggesting that *mutY* deletion or complementation with variant did not confer advantage (Fig S9c). These results suggest that the absence of prior antibiotic selection deletion or the presence of *mutY* variant does not confer an advantage (Fig S9c-d).

To test these conclusions, we performed the co-infection experiment with the strains that were subjected to three rounds of selection in the host in the presence or absence of antibiotics as described in Fig 4. Peritoneal macrophages were infected with a combination *Rv+Rv*Δ*mutY* or *Rv+Rv*Δ*mutY::mutY* or *Rv+Rv*Δ*mutY::mutY-*R262Q (Fig 5a). 24 h p.i cells were either treated or not treated with an antibiotic for the subsequent 72 h, and total CFUs were enumerated as described above to evaluate the survival rates. As expected, there was no difference in the CFUs either at 4 h p.i (Fig 5b-e) or 24 h p.i (data not shown). At 96 h p.i *Rv*Δ*mutY* and *Rv*Δ*mutY::mutY-*R262Q strains showed a distinct advantage over *Rv* both in the absence or presence of antibiotics. Importantly, *Rv*Δ*mutY::mutY* did not show any real advantage over *Rv* under any conditions. These results suggest that subjecting deletion or variant strains repeatedly to antibiotic stress in the host helps evolve strains that can outcompete the wild-type strain.

**Figure 5.**
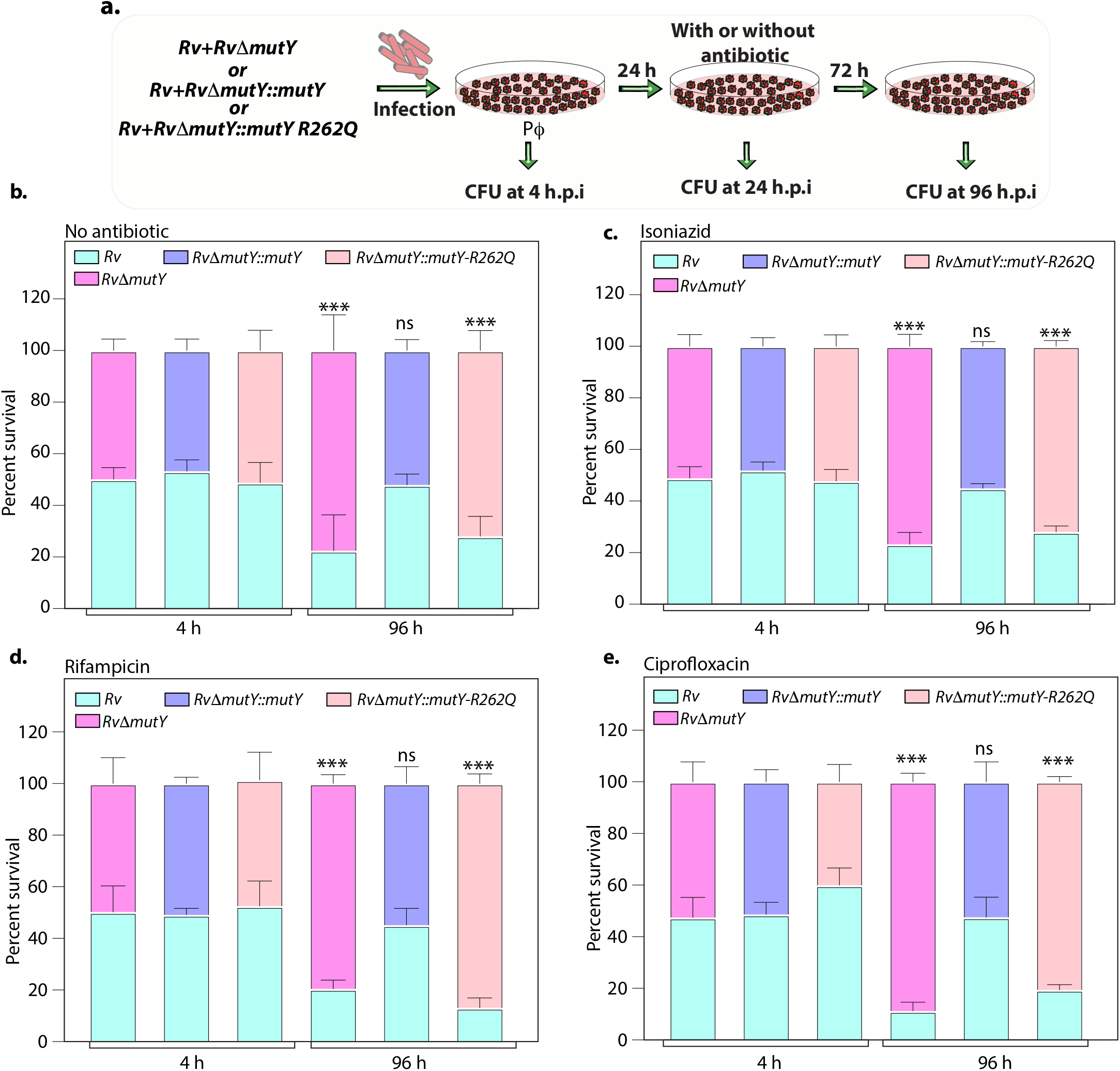
Variant of mutY outcompete Rv in competition experiment. **a)** Schematic representing the competition experiment performed in peritoneal macrophages. Strains that were obtained after three rounds of infection in the peritoneal macrophages were used to perform a competition experiment (Fig 4a). **b-e)** Percent survival of *Rv, Rv*Δ*mutY, Rv*Δ*mutY::mutY, Rv*Δ*mutY::mutY R262Q* in the absence and presence of antibiotics. Two biologically independent experiments with each experiment performed in technical triplicates. Data represents one of the two biological experiments. Data represents mean and standard deviation. Statistical analysis (two-way ANOVA) was performed using Graph pad prism software. *** p<0.0001, **p<0.001 and *p<0.01.

### *MutY* variant exhibits enhanced survival *in vivo*

Upon entering the host macrophages, *Mtb* encounters multiple stresses that impede its growth. To survive and grow in such a hostile environment, *Mtb* employs multiple defense mechanisms (34). The treatment regime with anti TB drugs imposes a supplementary layer of stress on the pathogen. An auxiliary mechanism used by the pathogen is to accumulate mutations in its genome that improve its ability to combat antibiotic and host-induced stress. We asked if the variant mutations identified in DNA repair genes are one such auxiliary mechanism? To test this hypothesis, we performed guinea pig infection experiments using *Rv, Rv*Δ*mutY*, *Rv*Δ*mutY::mutY,* and *Rv*Δ*mutY::mutY-*R262Q (Fig 6a). CFUs were enumerated after 1 and 56 days post-infection. CFUs obtained on day 1 showed the deposition was equal for wild type, mutant, and completed strains. Gross and histopathology analysis of infected lungs showed the presence of well-formed granulomas (Fig 6b-c). Significantly, 56 p.i, *Rv*Δ*mutY, Rv*Δ*mutY::mutY-*R262Q strains showed ∼5 fold superior survival than *Rv Rv*Δ*mutY::mutY,* suggesting that the variant mutant identified indeed confers advantage (Fig 6d).

**Figure 6.**
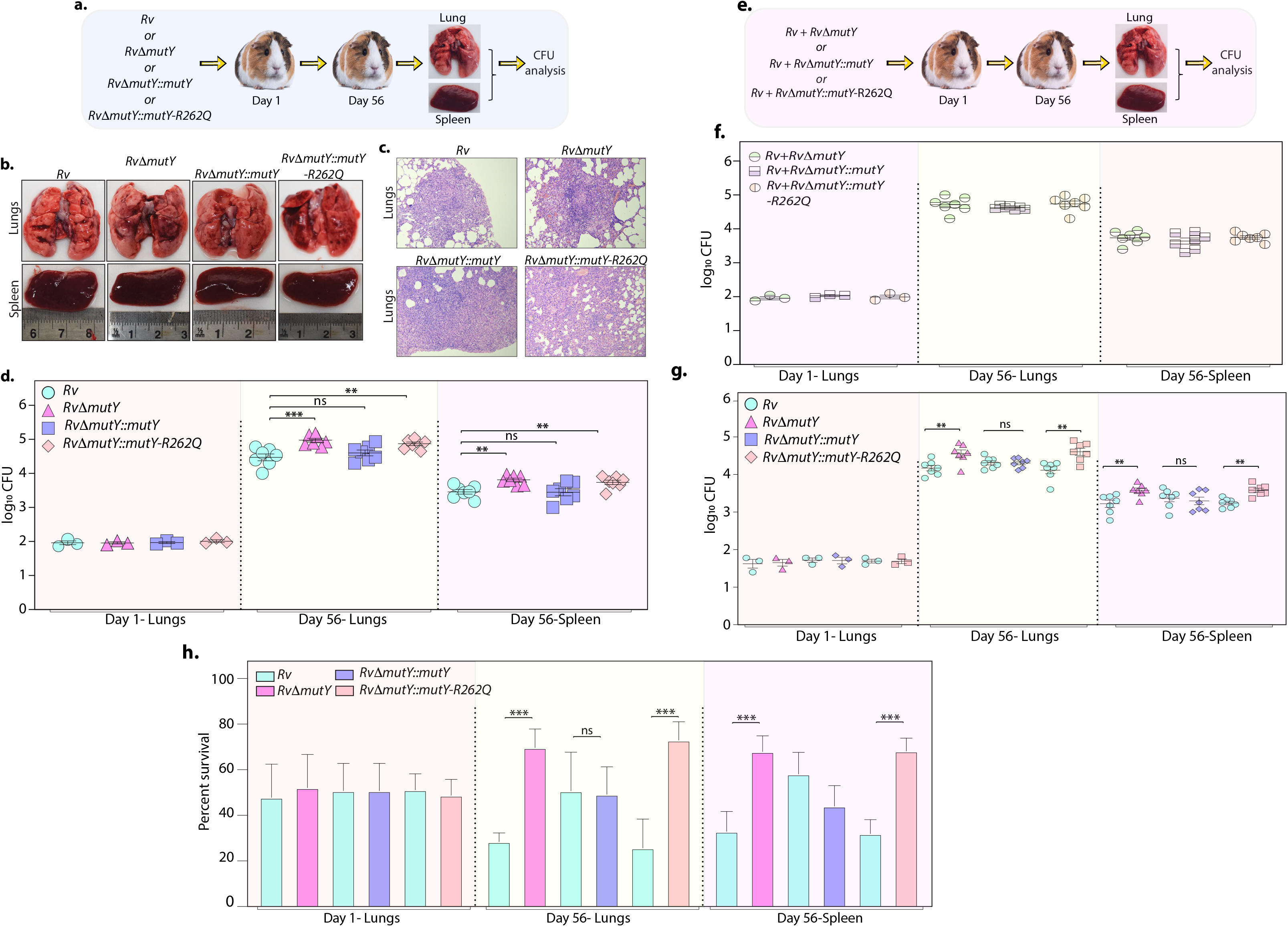
Perturbation of DNA repair results in enhanced survival in vivo. **a)** A schematic representation of guinea pig infection experiment. **b)** Gross histopathology of lungs and spleen of infected guinea pigs. **c)** Haematoxylin and eosin staining of infected lungs tissue showing the well-formed granuloma. Magnification 10X. **d)** Guinea pigs were challenged with *Rv, Rv*Δ*mutY, Rv*Δ*mutY::mutY, Rv*Δ*mutY::mutY-R262Q* via aerosol route. CFUs were enumerated at day 1 post-infection for determining the initial load in the lungs (n=3). The CFUs at one day p.i are represented for the whole lung. At 56-day p.i, lungs and spleen were isolated for determining the survival (CFU / ml, n=7). **e)** Outline showing the competition experiment performed in guinea pigs. **g)** CFU enumeration of *Rv + Rv*Δ*mutY*, *Rv + Rv*Δ*mutY::mutY* and *Rv + Rv*Δ*mutY::mutY-*R262Q in the lungs and spleen of guinea pigs at one day (whole lung, n=3) and 56-days p.i (CFU/ml, n=7). Statistical analysis was performed using two-way ANOVA. Graphpad prism was employed for performing statistical analysis. *** p<0.0001, **p<0.001 and *p<0.01. **h)** Survival of each strain at indicated time points in the mixed infection. **i)** Percent survival at 1- and 56-days in the competition experiment. Statistical analysis was performed using Unpaired t-test. *** p<0.0001, **p<0.001 and *p<0.01.

Next, we determined the survival ability of *Rv*Δ*mutY, Rv*Δ*mutY::mutY*, and *Rv*Δ*mutY::mutY-*R262Q, when competed against wild type *Rv* strain (Fig 6e). The lung and spleen homogenates were plated on 7H11 to determine total CFUs (Fig 6f, Fig S10). The total CFUs were found to be comparable in both lungs and spleen across all combinations (Fig 6f). Lung and spleen homogenates were plated on either Kanamycin (*Rv*) or Hygromycin (*Rv*Δ*mutY, Rv*Δ*mutY::mutY* or *Rv*Δ*mutY::mutY-*R262Q) containing plates to determine survival (Fig 6g). As was the case with independent infections, *Rv*Δ*mutY,* or *Rv*Δ*mutY::mutY-*R262Q, showed 5 fold elevated CFUs compared with *Rv* (Fig 6g). When plotted as percent survival, the data clearly showed that *mutY* deletion or complementation with the variant confers a survival advantage to the pathogen (Fig 6h). Collectively, results show that variant mutations identified in DNA repair genes using GWAS, abrogate their function and contribute to better survival under antibiotic/host stress conditions.

## Discussion

The whole-genome sequence (WGS) of clinical strains provides vital information about many aspects such as; drug resistance, the transmission of resistance, acquisition of resistance mutations, the evolution of compensatory mutations, and evolution of resistance in patients (35). A recent WGS analysis of 10219 diverse *Mtb* isolates successfully predicted mutations associated with pyrazinamide resistance (36). Large-scale GWAS, employed initially for analyzing human genome data, is an invaluable tool to delineate the mutations that confer antibiotic resistance in bacteria (37). Using the PhyC test on 116 clinical strains of *Mtb*, *ponA1* was identified as one of the targets of independent mutation (9). Analysis of 161 *Mtb* genomes identified polymorphisms in the intergenic regions that confer resistance to *p*-aminosalicylic acid (PAS) through overexpression of *thyA* and *thyX* (11). GWAS of 498 sequences revealed that mutation in alanine dehydrogenase correlated with the resistance to a second-line drug D-cycloserine (38). Evaluation of 549 strains led to the identification of *prpR*, which in an *ex vivo* infection model confers conditional drug tolerance through regulation of propionate metabolism (10). Largest GWAS involving 6465 clinical isolates uncovered novel resistance-associated mutations in *ethA* and *thyX* promoter, associated with ethionamide and PAS resistance, respectively (23).

We performed a gene-based GWAS analysis on a large dataset of susceptible and MDR/XDR clinical strains (Fig 1). We identified mutations in the known direct targets of both first and second-line antibiotics and few recently reported genetic variants using association analysis (Table 1-3). Besides, analysis captured mutations in genes that are involved in cell metabolism (Fig 2). This finding supports a recent study in *E.coli,* where drug treatment led to the acquisition of mutations in the metabolic genes that impart drug resistance (29). We speculate that these mutations may be compensatory mutations that placate the fitness cost associated with antibiotic resistance and hence might be the consequence of antibiotic resistance.

Most importantly, for the first time, we identified a significant association between mutations in three DNA repair pathway (BER, NER, and HR) genes, namely *mutY, uvrA, uvrB,* and *recF*, with MDR and XDR-TB phenotype. This study is in line with the other pathogens *Pseudomonas aeruginosa*, *Helicobacter pylori*, *Neisseria meningitides,* and *Salmonella typhimurium*, where mutations or deletions in the DNA repair genes were identified in the clinical isolates (39). Moreover, the deletion of *ung* and *udgB*, BER pathway genes, independently or together, provides a survival advantage to the bacteria (40). It is known that lineage 2 clinical strains have polymorphisms in the DNA repair and replication genes (41, 42). However, we identified mutations in DNA repair genes in lineage 4, suggesting that the phenomenon is not confined to lineage 2 (Fig S8a). Mutations in the *dnaA,* DNA binding protein that initiates replication, confers a survival advantage in the presence of isoniazid (43). We also identified a mutation in the *dnaA*, which is associated with drug resistance (Table 2).

We investigated if the mutations in DNA repair pathways are the cause or consequence of antibiotic resistance. We functionally validated the identified mutations of two different pathway genes, *mutY* and *uvrB*, in *Msm* using gene replacement mutants. Mutation frequency analysis suggests that GWAS identified mutations Arg262Gln and Ala524Val, in *mutY* and *uvrB*, respectively, abrogated their function (Fig 3). This agrees with the previously published study, wherein WGS analysis of antibiotic susceptible strain isolated from a patient showed a non-synonymous mutation in UvrB (A582V). Notably, the variant strain eventually evolved into XDR-TB over 3.5 years after the first and second-line drugs treatment (44). Therefore, we propose that mutations identified in DNA repair genes contribute to the evolution of antibiotic resistance (Fig S6).

Poor adherence of the patients to the antibiotic regimen is the leading cause of the emergence of drug resistance. The continual treatment of antibiotics provides sufficient time to evolve strains into MDR or XDR. We emulated the condition by repeatedly exposing strains to antibiotics in the *ex vivo* model, followed by growing them *in vitro* without antibiotics. This led to the improved survival of *Rv*Δ*mutY* and *Rv*Δ*mutY::mutY-*R262Q (Fig 4). We determined the survival of *ex vivo* evolved strains in the mixed infection scenario. Competition experiment using the evolved strains showed that *Rv*Δ*mutY* and *Rv*Δ*mutY::mutY-*R262Q could outcompete *Rv* (Fig 5). Host stress drives the selection of the bacterial population that acquires the ability to withstand adverse conditions. We evaluated the survival of the different strains in guinea pigs. Results suggest that *Rv*Δ*mutY* and *Rv*Δ*mutY::mutY-*R262Q exhibit improved survival. Similarly, we observed that *Rv*Δ*mutY* and *Rv*Δ*mutY::mutY-*R262Q could successfully outcompete *Rv* in the guinea pig model of infection (Fig 6).

Using the GWAS approach and functional validation of the clinical mutations identified in the BER and NER pathway, we established a novel link between compromised DNA repair and the evolution of antibiotic resistance (Fig 7). Collectively, the data presented here for the first time suggests that the evolution of MDR or XDR-TB is a consequence of the loss of function of DNA repair genes. The presence of mutations in DNA repair genes can be an early stage diagnostic marker for the evolution of the strain into MDR/XDR-TB. Molecular diagnosis of DNA repair gene mutations at the onset of infection should help design better therapies to impede the evolution of these strains into MDR or XDR. These findings indicate that compromised DNA repair in bacteria can contribute to the accumulation of mutations, which can evolve into MDR and XDR-TB with continued antibiotics treatment and selection.

**Figure 7.**
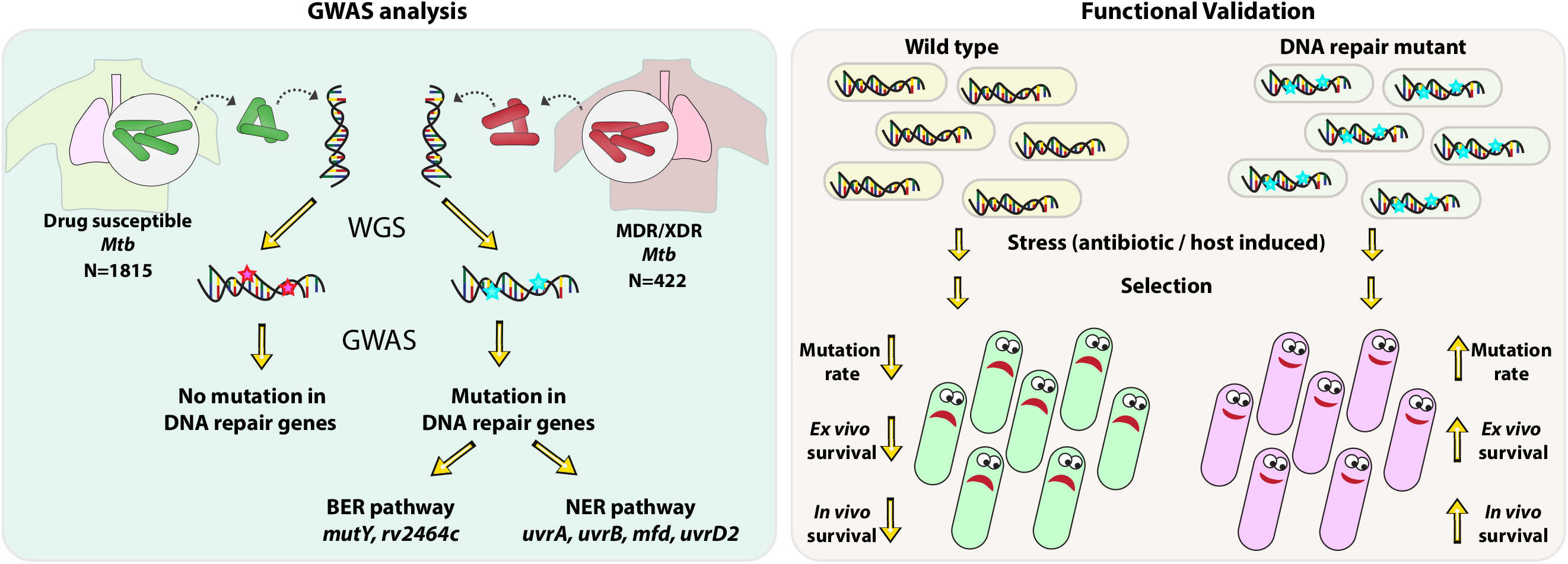
Model. Model depicts the analysis and subsequent validation. GWAS revealed mutation in three DNA repair pathway genes in MDR/XDR strains. Based on GWAS, we proposed that mutations in DNA repair genes contribute towards the evolution of antibiotic resistance in *Mtb*. Functional validation was performed using the gene replacement mutants of BER and NER pathway genes in *msm* and *Mtb*. *In vitro*, *ex vivo* and *in vivo* experiments show that compromised DNA repair pathway leads to evolution of MDR/XDR-TB.

## Materials and Methods

### Sequence retrieval and variant calling

Accession numbers for 2773 clinical strains of *Mtb* were obtained from 10 previous studies (Table S1), representing all four lineages from 9 countries. Sequence data was retrieved from EBA (https://www.ebi.ac.uk/) and NCBI database (https://www.ncbi.nlm.nih.gov/), and quality filtered using Trimmometic software (45). Adapter and *E. coli* sequence contaminations were removed, and the sequences analyzed at a sliding window of 4 bp, and those with an average phread value of 15 were clipped. Parameters for the Trimmometic based QC was set as (Leading:3, slidingwindow:4:15 trailing:3 minlen:60). Reads with a filtered length of <60 bp were removed. The filtered SE/PE reads were mapped on the H37Rv reference genome (Accession number –ASM19595) using BWA mem algorithm (46). GATK pipeline was used for sorting, PCR duplicates removal and realignment of the sequences (47). Variants (SNP and small InDels) were predicted in a batch mode with Platypus script (48). SNPs with read depth <5 or mapping quality of <20 were marked as missing. SNPs with missing calls in >40% of the accessions were removed from the analysis. Finally,∼160,000 SNPs from 2,773 accessions having a variant dataset with MAF of >1% were selected for the final analysis. The .vcf file was converted to .hmp file with Tassel (49).

### Association analysis

Genome-wide association analysis was carried out with 1,815 susceptible strains and 422 MDR/XDR strains (Table S2). A genome association and prediction integrated tool (GAPIT) based on a compressed mix linear model was performed under the R environment (50). VanRaden algorithm was employed for calculation of Kinship matrix. Kinship matrix assessed the relatedness among the strains included in the association panel. Principal components in GAPIT were employed classify the population structure. An association mapping analysis was carried out by combining the population structure analysis and the relative kinship matrix. P-values, R2, and marker effect values were obtained from the association mapping analysis. The false discovery rate (FDR) adjusted p-values in the GAPIT software is highly stringent as it corrects the effects of each marker based on the population structure (Q) as well as kinship (K) values and often leads to overcorrection (21, 22). We selected a corrected p-value of 10^-5^ as the threshold for selecting associated genes. The associated SNPs were annotated using the snpEff v4.11 (51). A snpeff database was generated using the H37Rv (ASM19595v2, was used as a reference for mapping). The corresponding .gff file and the SNPs were annotated based on their position on the genome. Dot-plot, Manhattan’s plot, and volcano plot were generated using custom R scripts.

### Analysis of Mutation frequency

Antibiotic sensitive cultures of *msm, msm*Δ*mutY, msm*Δ*mutY::mutY, msm*Δ*mutY::mutY- R262Q*, *msm*Δ*uvrB, msm*Δ*uvrB::uvrB* and *msm*Δ*uvrB::uvrB-A524V* were grown in 7H9-ADC medium A_600_ ∼0.6 and 50,000 cells/ml were inoculated in fresh 10 ml medium. Cultures were grown for six days in a 37^0^C incubator at 200 pm. On the 15^th^ day, appropriate dilutions were plated on 7H11-OADC plain plates for determining the load, and 1ml was plated on rifampicin (50 μg/ml). *Rv, Rv*Δ*mutY*, *Rv*Δ*mutY::mutY* and *Rv*Δ*mutY::mutY* R262Q strains were inoculated at A_600_ *∼*0.1. Strains were grown to A_600_ *∼*0.6 and plated on 7H11-plain or rifampicin (2 μg/ml) or isoniazid (10 μg/ml) plates to calculate the mutation frequency. Two biologically independent experiments were performed for determining mutation frequency. Each experiment was performed in technical triplicates. Data represent one of the two biological experiments. Data represent mean and standard deviation. Statistical analysis (two-way ANOVA) was performed using Graphpad prism software. *** p<0.0001, **p<0.001 and *p<0.01.

### Survival of strains *ex vivo*

Balb/c mice were injected with thioglycollate, and 72 h post injection, peritoneal macrophages were isolated. One million cells were seeded in each well of a 6 well plate. Cells were infected with *Rv, Rv*Δ*mutY*, *Rv*Δ*mutY::mutY,* and *Rv*Δ*mutY::mutY* R262Q independently at an MOI of 1:5. After 24 h p.i cells were treated with rifampicin (1μg/ml), isoniazid (1μg/ml) and ciprofloxacin (2.5 μg/ml). Cells were lysed at 120 h p.i using 0.05% SDS, and the bacteria was extracted. Bacteria was washed thrice with PBS to ensure the removal of SDS. Extraced bacteria were cultured in 7H9-ADC medium without antibiotics for 5-7 days up to A_600_∼0.4. These culture were used for the next round of infection. The whole process was repeated three times. During the fourth round of infection, CFUs were enumerated at 4 h p.i and 96 h p.i to determine different strains’ survival. Percent survival was calculated by normalized CFU obtained at 96 h with respect to the respective CFUs obtained at 4 h. Two biologically independent experiments were performed. Each experiment was performed in technical triplicates. A representative experiment is shown in Fig 4.

Similarly, the strains obtained after three rounds of infection from the above experiment were used for the competition experiment. *Rv+Rv*Δ*mutY*, *Rv+Rv*Δ*mutY::mutY,* and *Rv+Rv*Δ*mutY::mutY*-R262Q were mixed in 1:1 ratio. 24 h p.i the cells were either not treated or treated with the antibiotics (isoniazid-1μg/ml, rifampicin- 1μg/ml and ciprofloxacin- 2.5 μg/ml) for subsequent 72 h. CFUs were enumerated at 4 h and 96 h p.i on 7H11-plain (all strains), 7H11-Kan (Rv), and 7H11-Hyg (*RvΔmutY, RvΔmutY::mutY*, and *RvΔmutY::mutY-R262Q*) plates. Percent survival was calculated as [CFUs on Kan or Hyg plates / (CFUs on Kan + CFUs on Hyg)]*100. Data represent one of the two biological experiments. Data represent mean and standard deviation. Statistical analysis (two-way ANOVA) was performed using Graph pad prism software. *** p<0.0001, **p<0.001 and *p<0.01.

### Guinea pig infection

*Rv, Rv*Δ*mutY*, *Rv*Δ*mutY::mutY* and *Rv*Δ*mutY::mutY-*R262Q cultures were grown up to A_600_ ∼0.8. For preparing single cell suspension, *Rv, Rv*Δ*mutY*, *Rv*Δ*mutY::mutY* and *Rv*Δ*mutY::mutY-*R262Q cultures were pelleted at 4000 rpm at room temperature (RT). After suspending cells in saline, cells were passed through a 26 ½ gauge needle to obtain single-cell suspension. 1 x 10^8^ cells were taken in the 15 ml saline for infection. Female outbred Hartley guinea pigs were challenged using a Madison chamber calibrated to deliver ∼ 100 bacilli/lung through aersolic route. For determining the deposition of bacteria in the lungs of guinea pigs (n=3) CFUs were enumerated at one day post-infection on 7H11-plain containing plates for each strain. Survival of each strain was determined at 56-day post-infection in the lungs and spleen (n=7). *Rv+Rv*Δ*mutY*, *Rv+Rv*Δ*mutY::mutY,* and *Rv+Rv*Δ*mutY::mutY* R262Q were mixed in 1:1 ratio, and the guinea pigs (n=10 per strain) were challenged as described above. CFUs were enumerated at 1- and 56 – days post-infection on 7H11-plain, Kanamycin or Hygromycin containing plates. Data represent Standard Error Mean (SEM). Statistical analysis (two-way ANOVA) was performed using Graph pad prism software. *** p<0.0001, **p<0.001 and *p<0.01. Percent survival was calculated as: [CFUs on Kan or Hyg plates / (CFUs on Kan + CFUs on Hyg)]*100. Data represents Standard deviation and mean. Statistical analysis (Unpaired t-test) was performed using Graph pad prism software. *** p<0.0001, **p<0.001 and *p<0.01. All guinea pig infection experiments were performed at the same time. Guinea pigs were not treated with any antibiotics before or after infection

## Acknowledgement

This work was funded by the Department of Biotechnology, Government of India (BT/PR13522/COE/34/27/2015), and J.C Bose fellowship (JCB/2019/000015). SN is an Senior Project Associate in the J.C Bose fellowship (JCB/2019/000015). We thank Tuberculosis Aerosol Challenge Facility at ICGEB animal facility and staff for their help in performing animal infection experiments. We are thankful to the bio-containment facility (BSL3) at NII. Vector-pYUB1471 is a kind gift from Prof. William R. Jacobs’s laboratory.

## Contribution

SN, KP, and VKN conceptualized the study. KP performed genome assembly and GWAS analysis. UV’s lab provided *msmΔmutY* and *msmΔuvrB* knockouts. SN performed all functional validation *in vitro*, *ex vivo*, *in vivo* infection experiments. SN and PS performed *ex vivo* and *in vivo* infection experiments. SK helped in generating the gene replacement mutant. SN, KP, and VKN generated figures. VKN, UV, and YS supervised the study. SN, VKN wrote the manuscript.

## Ethics

Animal experiments protocol was approved by the Animal Ethics Committee of the National Institute of Immunology, New Delhi, India. The approval (IAEC#409/16) is as per the guidelines issued by the Committee for the Purpose of Control and Supervision of Experiments on Animals (CPCSEA), Government of India.

## Supplementary Material

### Methods

#### Generation of gene replacement mutant in *Mtb*

Upstream (5’ flank) and downstream region (3’ flank) of the *mutY* was amplified using the *Rv* genomic DNA. Flanks were digested with appropriate restriction enzyme and ligated with oriE+lambda cos and hygromycin resistance cassette (52). The allelic exchange substrate was digested to generate linearized substrate and electroporated in the recombineering proficient *Rv* strain harbouring pNit-ET plasmid (53). Colonies were screened post 3 weeks electroporation for gene replacement mutant.

#### Generation of complementation constructs

Wild type allele of *mutY* or *uvrB* was PCR amplified using *Rv* genomic DNA. The PCR product and the vector pSTL-giles was digested with NdeI and HindIII to generate pSTL- giles-*mutY* or pSTL-giles-*uvrB*. Subsequently the PCR product was ligated with the vector pSTL-giles. SapI based cloning was employed for the generation of pSTL-giles-*mutY- R262Q* or pSTL-giles-*uvrB-A524V.* Constructs pSTL-giles-*mutY* and pSTL-giles-*mutY- R262Q* were electroporated in the *msm*Δ*mutY* or *Rv*Δ*mutY* to generate *msm*Δ*mutY::mutY* and *msm*Δ*mutY::mutY-R262Q* or *Rv*Δ*mutY::mutY* and *Rv*Δ*mutY::mutY-R262Q.* Constructs, pSTL-giles-*uvrB* or pSTL-giles-*uvrB-A524V* were electroporated in *msm*Δ*uvrB* to generate *msm*Δ*uvrB::uvrB* or *msm*Δ*uvrB::uvrB- A524V*.

## Supplementary Tables

**Table S1:** Total number of clinical strains used in this study. The table contains the total number of clinical strains obtained from different studies.

**Table S2:** Clinical strains used for the Genome wide association analysis. The table contains the clinical strains which are used for performing GWAS analysis.

**Table S3:** Synonymous change identified in the association analysis. The table contains synonymous changes that are identified in the MDR/XDR strains.

**Table S4:** Non-synonymous change identified in the association analysis. The table contains non-synonymous changes that are identified in the MDR/XDR strains.

**Table S5:** Upstream gene variants identified in the association analysis. The table contains non-upstream gene variants identified in the MDR/XDR strains.

**Table S6:** Stop codon or frame shift mutations identified in the association analysis. The table contains Stop codon or frame shift mutations identified in the MDR/XDR strains.

**Table S7:** Codon usage of the strains in the MDR/XDR strains. The table contains codon usage in the MDR/XDR and *Rv* strain.

**Table S8:** Oligonucleotide used in the study. The table consist of oligonucleotide that are used in the study.

## Supplementary figure legend

**Figure S1.**
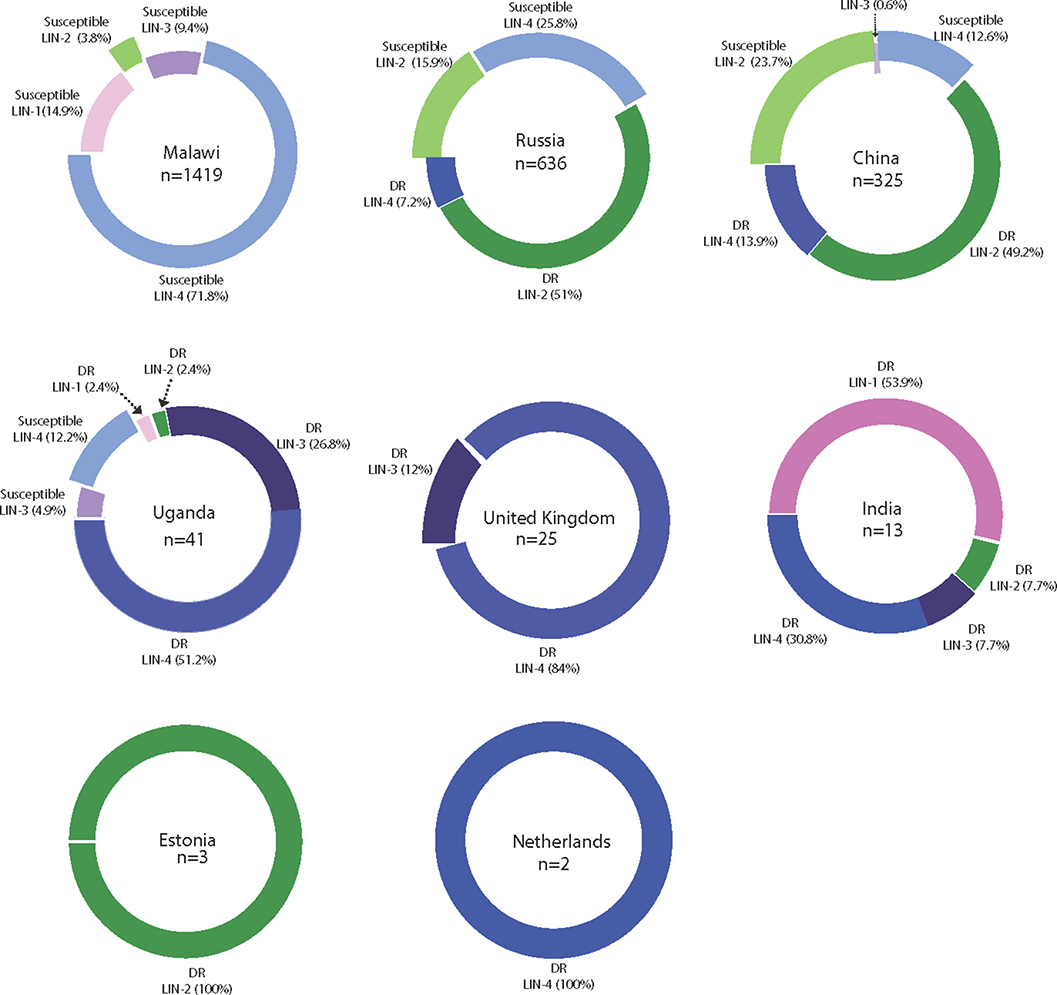
Country wide distribution of clinical strains. Each donut plot represents proportion of clinical strains used for the GWAS. Susceptible refers to the antibiotic sensitive strains. DR refers to ‘Drug Resistance’ to first- and second-line antibiotics.

**Figure S2.**
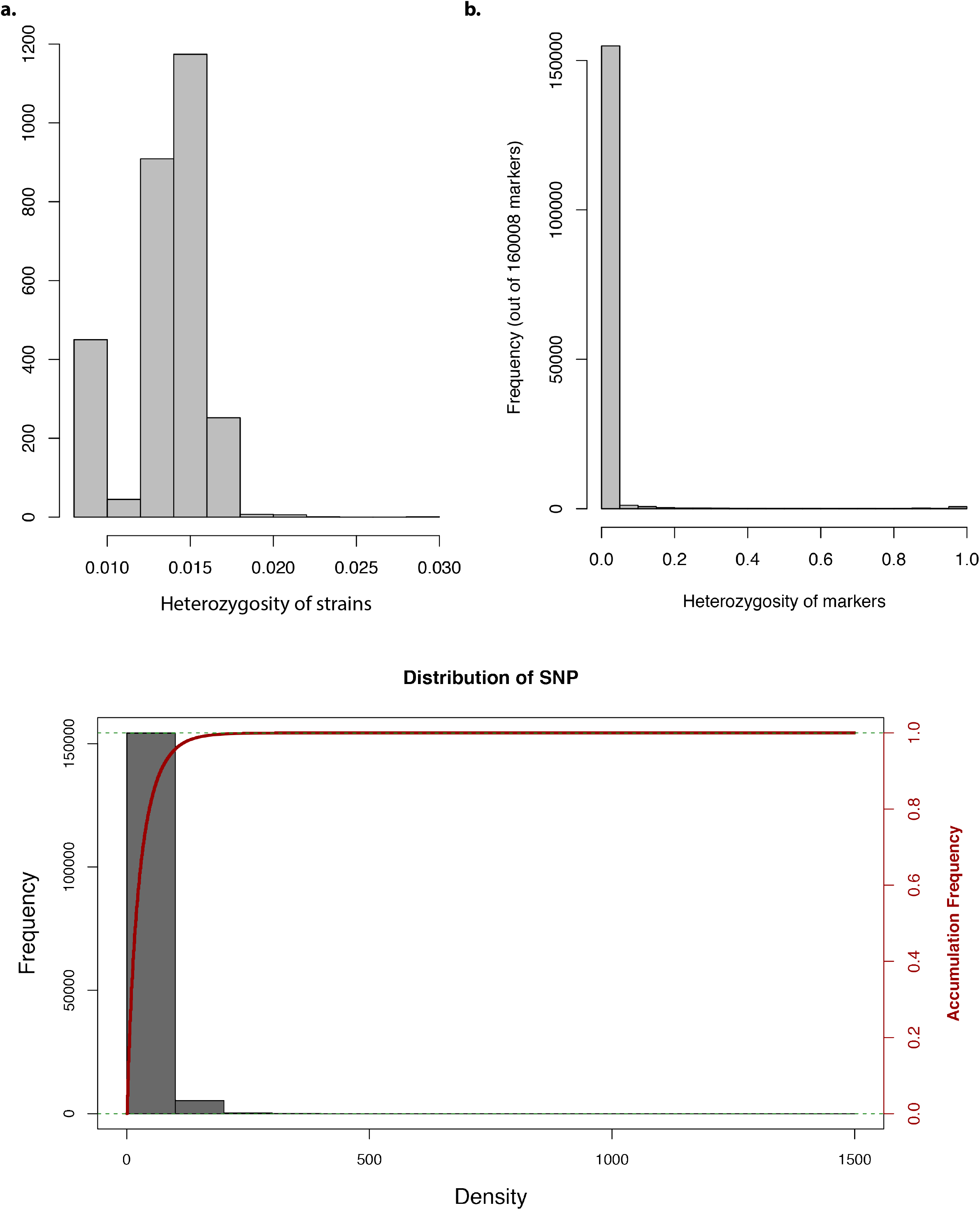
GWAS analysis. Bar graph representing the **a)** heterozygosity of strains, **b)** heterozygosity of markers, **c)** Frequency and accumulative frequency of marker density.

**Figure S3.**
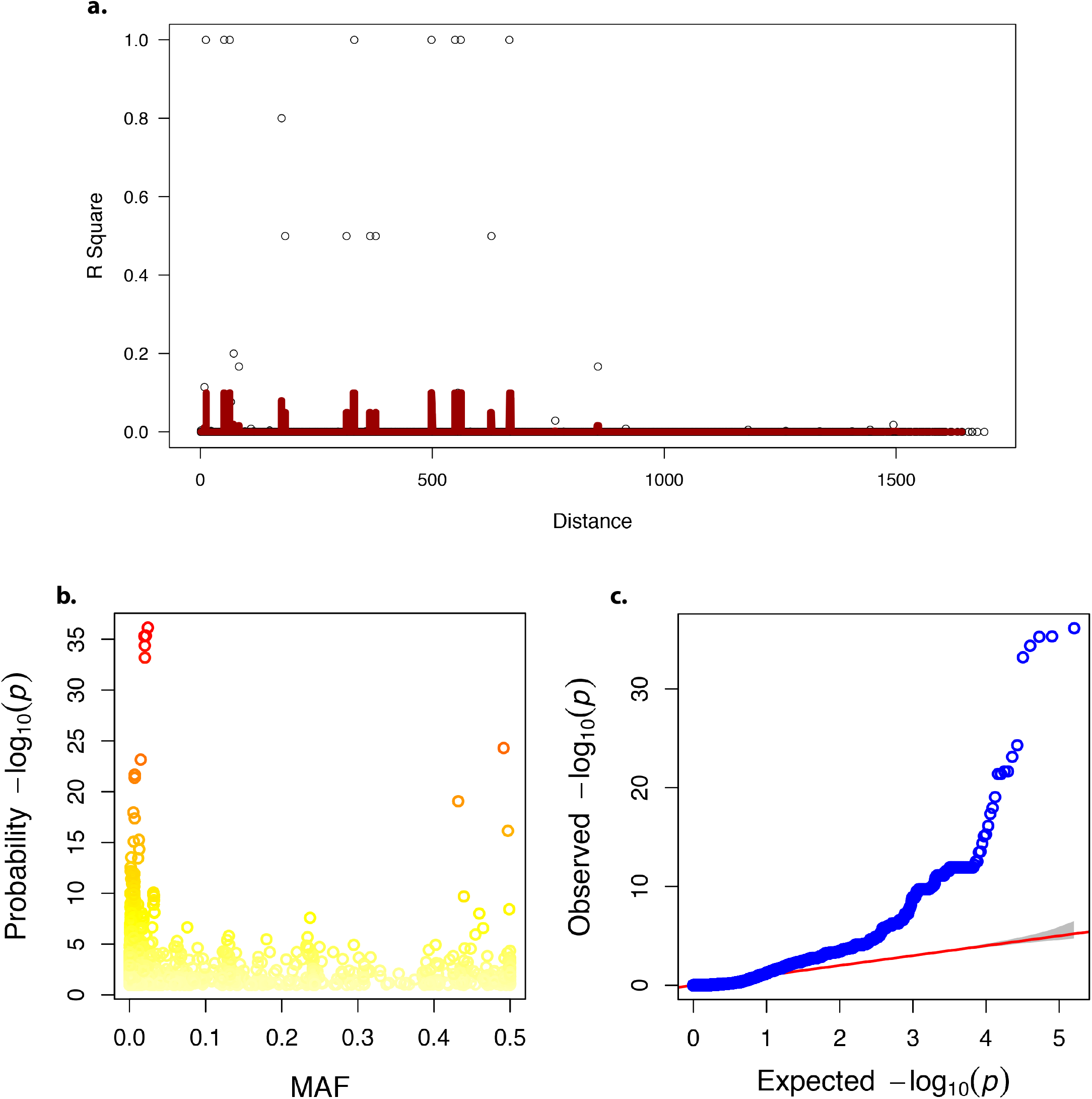
**a)** Linkage disequilibrium (LD) decay over distance. **b)** Minor Allele Frequency (MAF). **c)** Quantile-quantile (QQ) –plot of p-values.

**Figure S4.**
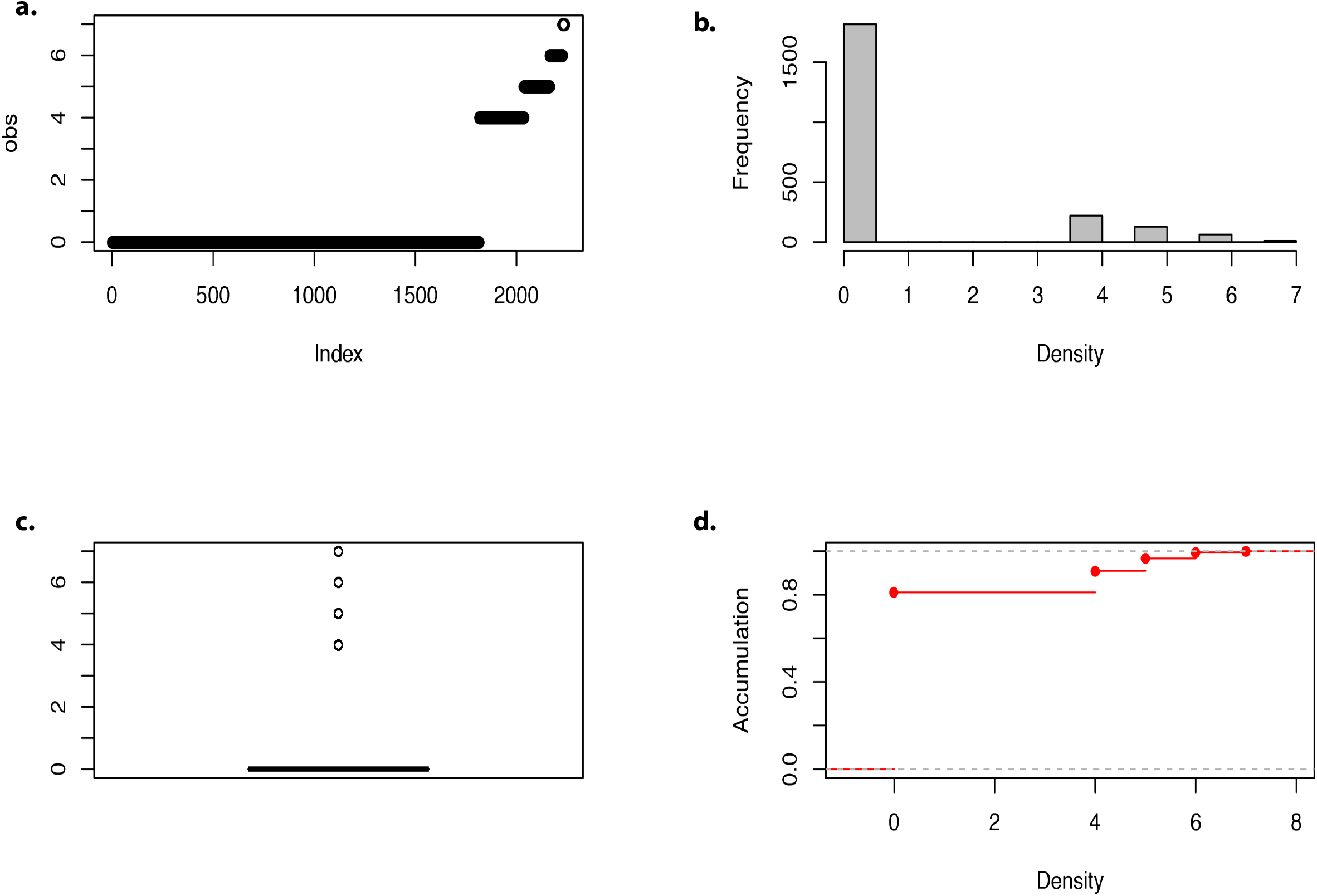
**a-d)** Density of markers.

**Figure S5.**
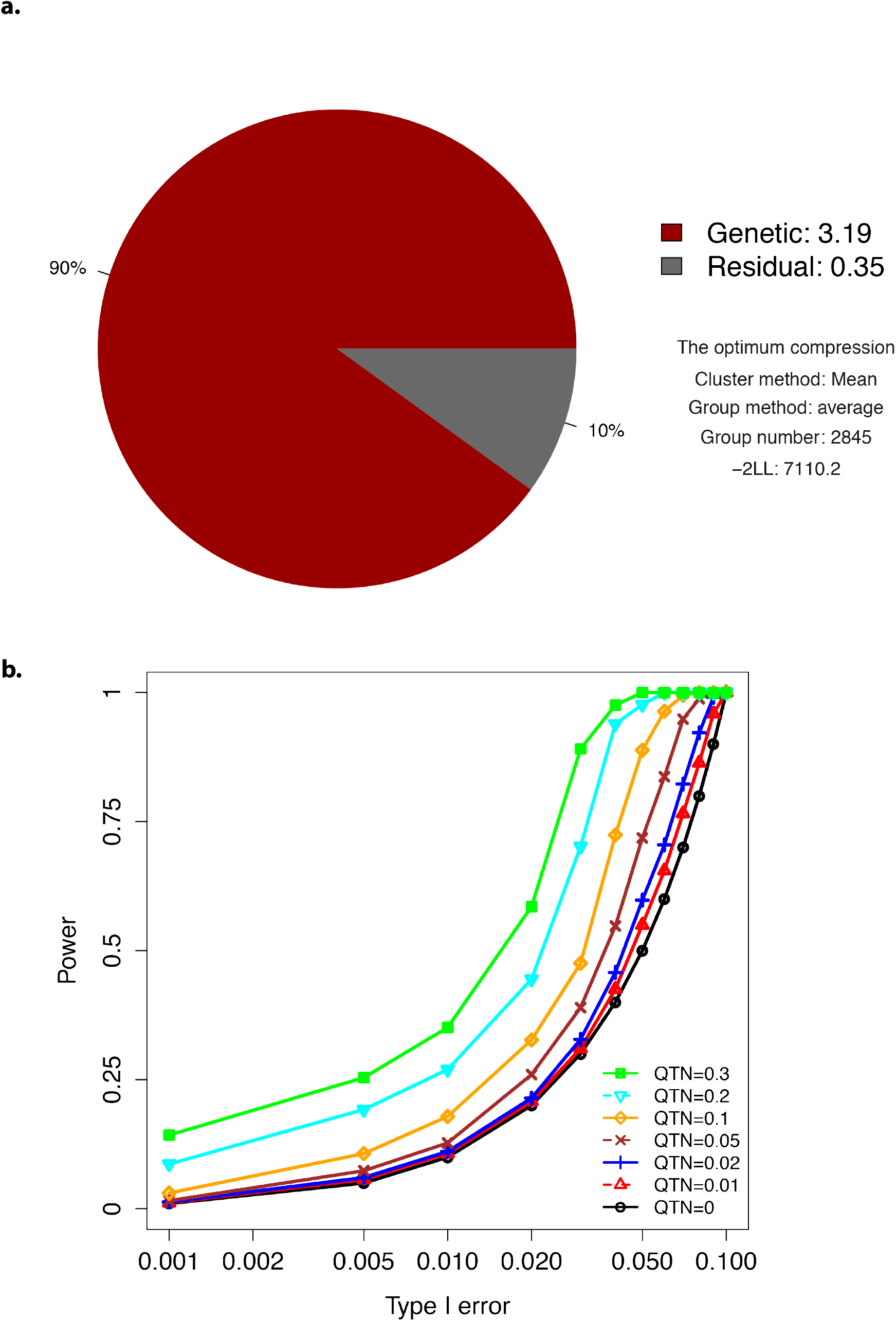
**a)** The profile for the optimum compression. **b)** Type-I error plot.

**Figure S6.**
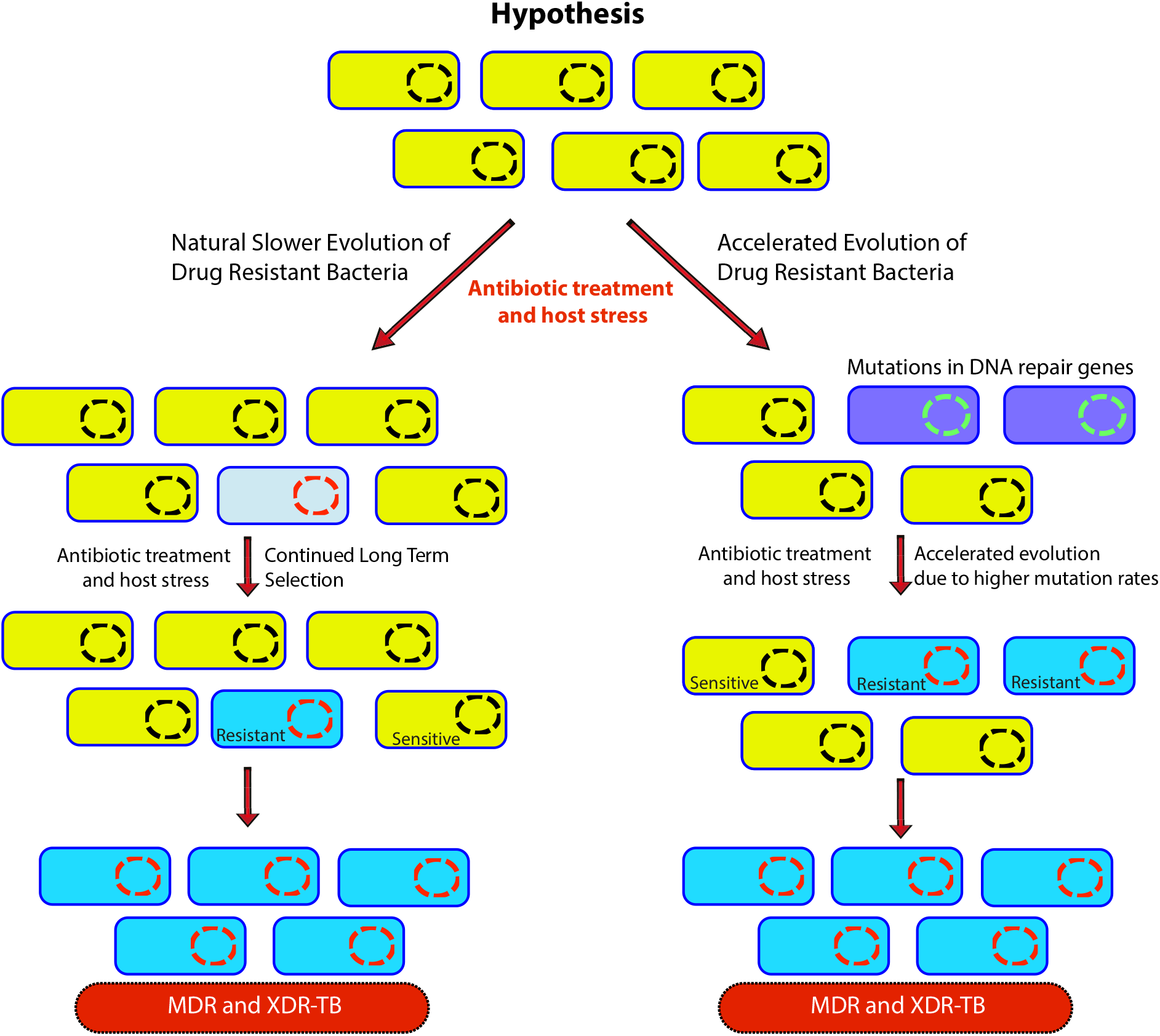
GWAS based hypothesis. In natural process of evolution host-imposed stress and antibiotic treatment results in the evolution of wild type bacteria in MDR/XDR in due course of time, however, in bacteria that possess compromised DNA repair pathway the evolution to MDR/XDR-TB is accelerated.

**Figure S7.**
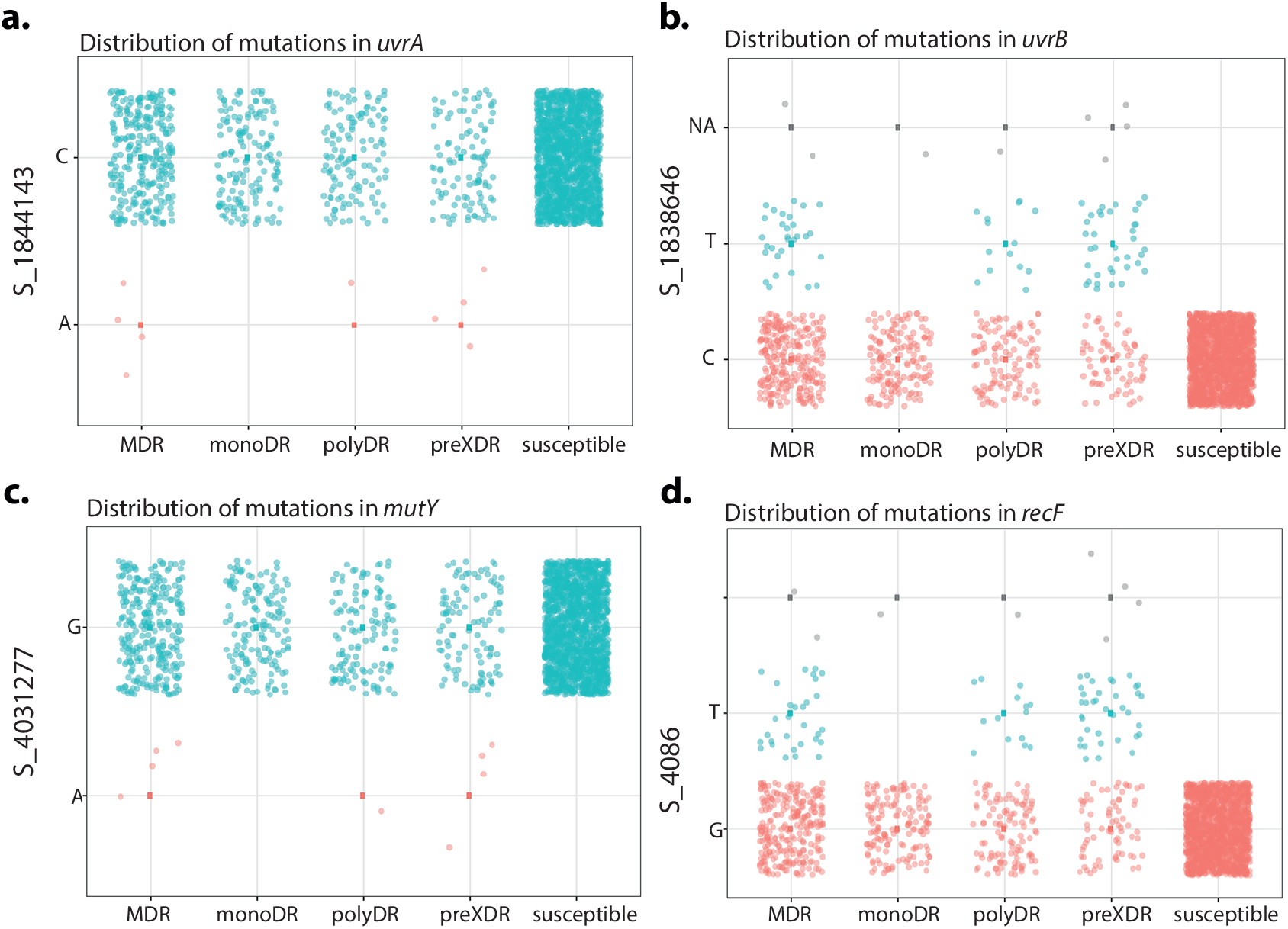
Distribution of mutations in the DNA repair genes. **a-d)** Distribution plot showing SNPs in *mutY, recF, uvrA* and *uvrB* identified in drug resistant strains. Wild type and the alternative alleles are shown.

**Figure S8.**
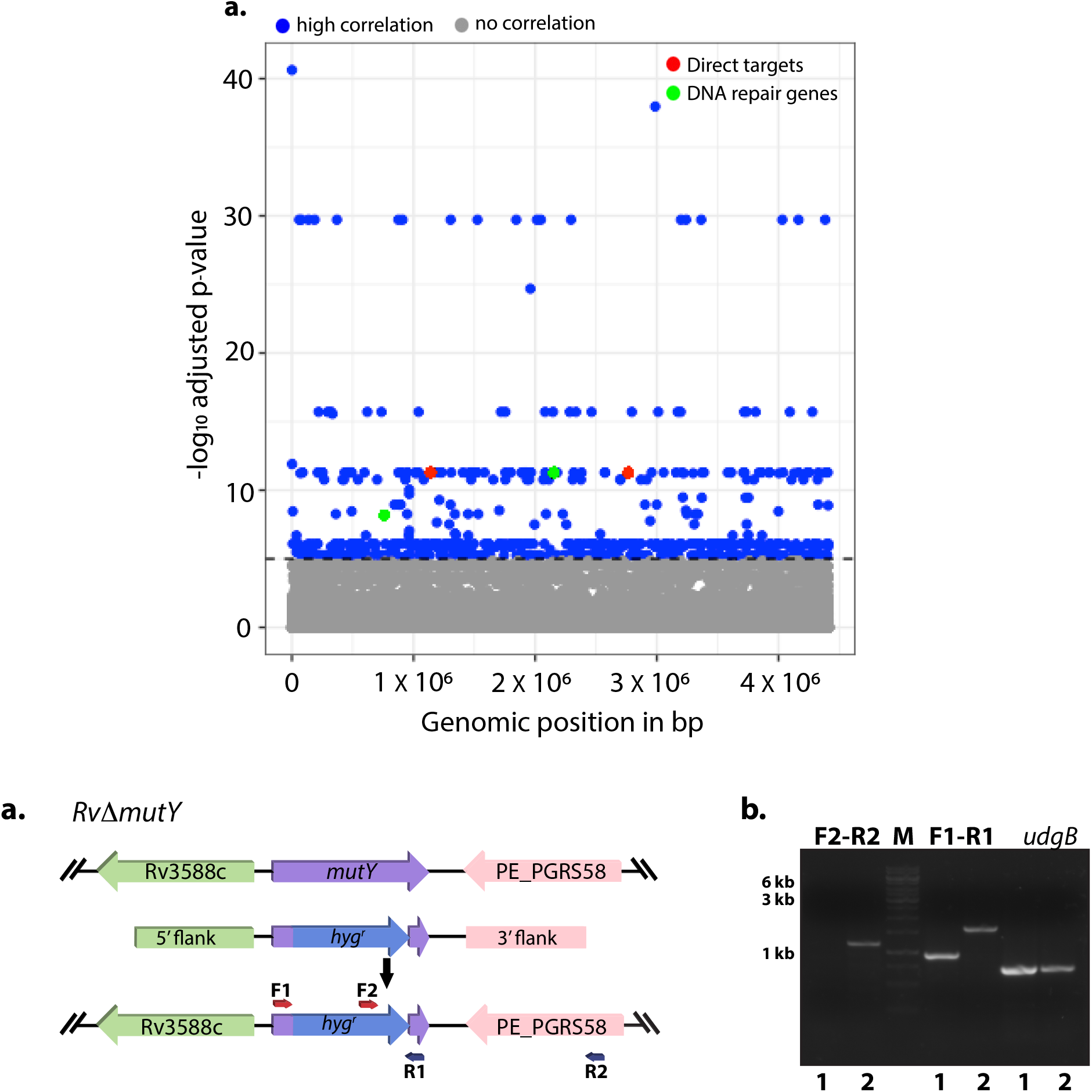
GWAS of lineage 4 strains identified mutations in the DNA repair genes. **a)** Manhattan plot showing the identification of genes that belongs to DNA repair and direct target of antibiotics. **b)** Schematic depicting the generation of gene replacement mutant of *mutY*. Native allele was disrupted by the *hygromycin^r^* cassette. b) PCR using F1-R1 (gene specific primers) resulted in the amplification of 900 bp in *Rv* and ∼1.6 kb in *Rv*Δ*mutY*. PCR using F2-R2 (*hygromycin^r^* cassette forward primer and reverse primer beyond the 3’ flank) resulted amplification in the *Rv*Δ*mutY* but not in *Rv*.

**Figure S9.**
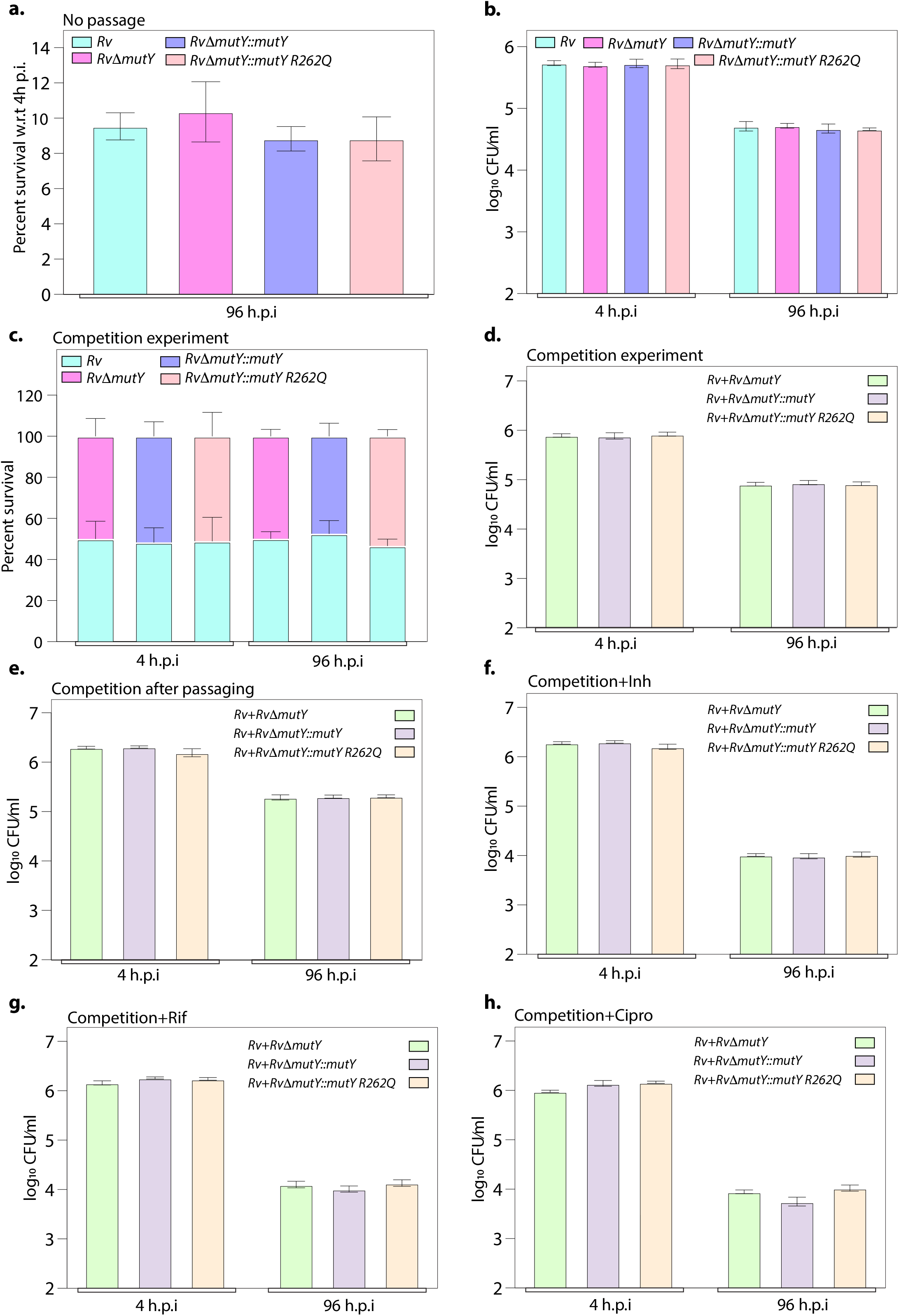
*Ex vivo* survival of strains. **a)** Survival of strains with respect to 4 h p.i in peritoneal macrophages. **b)** CFU enumeration of strains at 4 and 96 h p.i. **c)** Percent survival of each strain in the competition experiment. **d)** CFU enumeration of strains at 4 and 96 h p.i. in competition experiment. **e)** Competition experiment after passaging of strains *ex vivo*. In the absence of antibiotic, the CFU enumeration was performed at 4 and 96 h p.i. CFU enumeration in the presence of **f)** Isoniazid, **g)** Rifampicin, and **h)** Ciprofloxacin at 4 and 96 h p.i.

**Figure S10.**
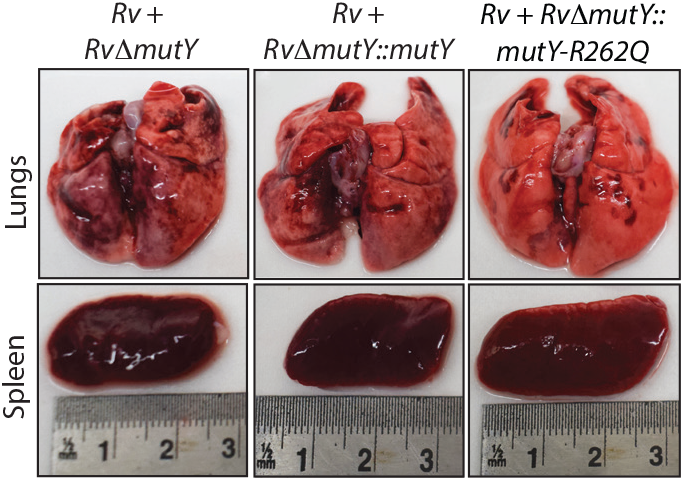
Gross histopathology of infected lungs and spleen isolated at 56-day post infection.

